# A virtual clinical trial of psychedelics to treat patients with disorders of consciousness

**DOI:** 10.1101/2024.08.16.608251

**Authors:** Naji L.N. Alnagger, Paolo Cardone, Charlotte Martial, Yonatan Sanz Perl, Iván Mindlin, Jacobo D Sitt, Leor Roseman, Robin Carhart-Harris, David Nutt, Pablo Mallaroni, Natasha Mason, Johannes G Ramaekers, Vincent Bonhomme, Steven Laureys, Gustavo Deco, Olivia Gosseries, Pablo Núñez, Jitka Annen

**Affiliations:** Coma Science Group, GIGA-Consciousness, University of Liège, Avenue de l’hôpital 11, Liège 4000, Belgium; Centre du Cerveau², University Hospital of Liège, Avenue de l’hôpital 11, Liège 4000, Belgium; Sorbonne Université, Institut du Cerveau - Paris Brain Institute - ICM, Inserm, CNRS, Paris 75013, France; Department of Information and Communication Technologies, Centre for Brain and Cognition, Computational Neuroscience Group, Universitat Pompeu Fabra, Barcelona, Spain; Department of Psychology, University of Exeter, Exeter, UK; Centre for Psychedelic Research, Department of Brain Sciences, Faculty of Medicine, Imperial College London, London, United Kingdom; Psychedelics Division, Neuroscape, University of California, San Francisco, San Francisco, CA, United States; Department of Neuropsychology and Psychopharmacology, Faculty of Psychology and Neuroscience, Maastricht University, Maastricht, The Netherlands; Anesthesia and Perioperative Neuroscience Laboratory, GIGA-Consciousness, GIGA Institute, University of Liège, Liège, Belgium; Department of Anaesthesia and Intensive Care Medicine, Liège University Hospital, Liège, Belgium; CERVO Brain Research Centre, Laval University, 2601 de la Canardière, Québec, G1J 2G3, Canada; Department of Data Analysis, University of Ghent, Henri Dunantlaan 1, Ghent 9000, Belgium

## Abstract

Disorders of consciousness (DoC), including the unresponsive wakefulness syndrome (UWS) and the minimally conscious state (MCS), have limited treatment options. Recent research suggests that psychedelic drugs, known for their complexity-enhancing properties, could be promising treatments for DoC. This study uses whole-brain computational models to explore this potential. We created individualised models for DoC patients, optimised with empirical fMRI and diffusion-weighted imaging (DWI) data, and simulated the administration of LSD and psilocybin. We used an in-silico perturbation protocol to distinguish between different states of consciousness, including DoC, anaesthesia, and the psychedelic state, and assess the dynamical stability of the brains of DoC patients pre- and post-psychedelic simulation. Our findings indicate that LSD and psilocybin shift DoC patients’ brains closer to criticality, with a greater effect in MCS patients. In UWS patients, the treatment response correlates with structural connectivity, while in MCS patients, it aligns with baseline functional connectivity. This virtual clinical trial lays a computational foundation for using psychedelics in DoC treatment and highlights the future role of computational modelling in drug discovery and personalised medicine.

## Introduction

The diverse phenomenological capacities of conscious experience are orchestrated by the emergence of spatiotemporally complex dynamics that facilitate the brain’s capacity to integrate information from differentiated sources^1–3^. These flexible dynamics are underpinned by two key aspects of the brain^4–6^: its physical wiring, represented by white matter structural connectivity obtained through diffusion weighted imaging (DWI), and neural function, approximated by the blood oxygen level dependent signal (BOLD) derived from functional magnetic resonance imaging (fMRI)^7^. Healthy brain dynamics are highly sensitive to perturbations and operate at a critical, or marginally subcritical regime^8–10^. Criticality refers to the state at the transition point between order and disorder. This behaviour enables efficient information processing by maximising adaptability, whilst preserving order^11–13^. Criticality is also associated with maximal sensitivity to perturbation and is thought to be one of the main dynamical mechanisms through which nature produces complexity^14^. Disruptions of brain function, in the presence of a healthy brain structure, can significantly alter conscious experience, such as during dreamless sleep or anaesthesia. Conversely, severe brain injury, whether due to anoxia or traumatic brain injury, causes major structural damage and subsequent functional deficits, which can result in coma and post comatose Disorders of Consciousness (DoC). In states of reduced consciousness such as DoC and anaesthesia from drugs like propofol and dexmedetomidine, the brain loses its capacity to integrate information, becomes more segregated, decreases in functional complexity, and shifts away from criticality^15–20^. DoC include unresponsive wakefulness syndrome (UWS) and minimally conscious state (MCS). UWS is characterised by wakefulness and reflex actions without awareness^21^, while MCS involves non-reflex behaviours (e.g., responses to command, visual pursuit) without functional communication^22^.

Classical psychedelics, such as lysergic acid diethylamide (LSD) and psilocybin, primarily act upon the 5-HT2A receptor and impart alterations in perception, mood and numerous cognitive processes^23^. Psychedelics have been proposed as promising treatments for various affective disorders in psychiatry^24–29^. Compared to normal wakefulness, the brain in the psychedelic state operates at even more unstable regime, at the edge of criticality, with a higher complexity^30–35^. Therefore, using psychedelics to push the brain dynamics of patients with DoC from a state of pathologically low complexity closer towards criticality, and more similar to the dynamics of a healthy brain, raises the possibility of a new treatment paradigm^36,37^. Such a treatment could acutely increase the conscious level of patients, potentially eliciting new conscious behaviours or enriching their phenomenological experience.

Administering psychedelics to patients with DoC presents significant challenges. Their legal status alone presents difficulties in obtaining licences for drug storage and management. Additionally, administering a substance that profoundly alters phenomenological experience to a population that cannot provide consent requires comprehensive ethical considerations^38^ and meticulous study design. Whole-brain computational modelling offers a way to circumvent these challenges by performing experiments *in-silico*^39^. Such investigations can also provide causal mechanistic insights into structural and functional dependencies supporting the brain dynamics in different states of consciousness^6,40–42^.

One effective way to study a dynamical system is to introduce a perturbation and observe its return to a baseline state. The perturbational complexity index (PCI) leverages this approach by calculating the algorithmic complexity of the resultant electroencephalography (EEG) signal following a transcranial magnetic current stimulation (TMS) pulse^43^. Similar, *in-silico* perturbation protocols using whole-brain models at the group level have been shown to distinguish between states of consciousness^30,42,44,45^. One of these approaches, the perturbative integration latency index (PILI), measures the amplitude and latency of the dynamical return to baseline from a modelled perturbation, thereby measuring brain dynamical stability. Increased sensitivity to a perturbation indicates a more unstable system, with dynamics that would be closer to criticality, consistent with a system with higher complexity.

The heterogeneity of functional deficiencies and structural lesions in patients with DoC makes personalised models for scientific investigation and medical applications an essential, yet aspirational target^46^. In this study, we created individualised generative whole-brain models based on Stuart-Landau oscillators, integrating the empirical BOLD fMRI data, and for the first time, personalised DWI tractography data from each patient with DoC. We calculated PILI from an *in-silico* perturbation protocol, to characterise the brain dynamics of states of consciousness including DoC, anaesthesia and the psychedelic state. We then utilised a framework incorporating empirical LSD and psilocybin fMRI data from healthy controls to virtually simulate drug administration on each individual patient with DoC, before assessing the resulting brain dynamics using PILI. By connecting physical reality with a digital representation, we aimed to create digital twins^47^ of patients with DoC, and enrol them onto a "phase zero" clinical trial to test the efficacy of this potentially paradigm-shifting treatment.

## Results

We built whole-brain computational models, optimised at the individual patient level to explore using psychedelic drugs as a treatment for DoC. We assessed brain dynamics using an *in-silico* perturbation protocol, which measures the return to baseline dynamics after a modelled perturbation. Firstly, we show how the response to perturbation can discriminate between different states of consciousness (i.e., DoC, anaesthesia from propofol, and dexmedetomidine, and the psychedelic state under LSD and psilocybin). Then, we simulated the administration of LSD and psilocybin separately upon individual patient models and assessed the modelled dynamics both at baseline and at the simulated psychedelic state. Lastly, we characterised the baseline predictive biomarkers of simulated treatment effect.

### Response to perturbation differentiates between states of consciousness

We first sought to establish an *in-silico* perturbational biomarker that can distinguish between states of consciousness, based on the level of consciousness, and phenomenological richness of experience. This could later serve as a proxy to assess the simulated treatment response in our individual patient models. To achieve this, we created Hopf whole-brain computational models optimised on the functional connectivity (FC) matrix obtained from the BOLD fMRI data corresponding to each state of consciousness, averaged at the group level (Figure 1). We used previously published data^48–52^ from several states of consciousness to create 13 different models: 3 for the DoC dataset (see Supplementary Table 1 for the demographics of the patients with DoC) (healthy controls (CNT) N=35, UWS N=20, and MCS N=26), 4 models based on healthy subjects under anaesthesia (propofol N=15, dexmedetomidine N=11, as well as a separate model of wakefulness for each dataset), and 4 models based on healthy subjects in the psychedelic state (LSD N=12, psilocybin N=12 and the two related placebo conditions). We also performed a supplementary analysis using a model built on another psilocybin dataset, presented in Supplementary Materials. Briefly, each model possesses two sets of parameters, the global coupling parameter (G), and the local bifurcation parameters (α). The global coupling parameter linearly scales the coupling between brain regions in the structural connectivity (SC) matrix. The local bifurcation parameters denote the dynamics of each node: negative values produce noisy activity, while positive values lead to stable oscillatory dynamics. After retrieving the optimal global coupling value, the local parameter optimisation was restricted to a nine-parameter space representing nine resting state networks (attention, auditory, default mode [DMN], frontoparietal, limbic, sensorimotor, precuneus, thalamus, visual) defined by an independent components analysis on the fMRI data from the healthy controls of the DoC dataset.

**Figure 1:**
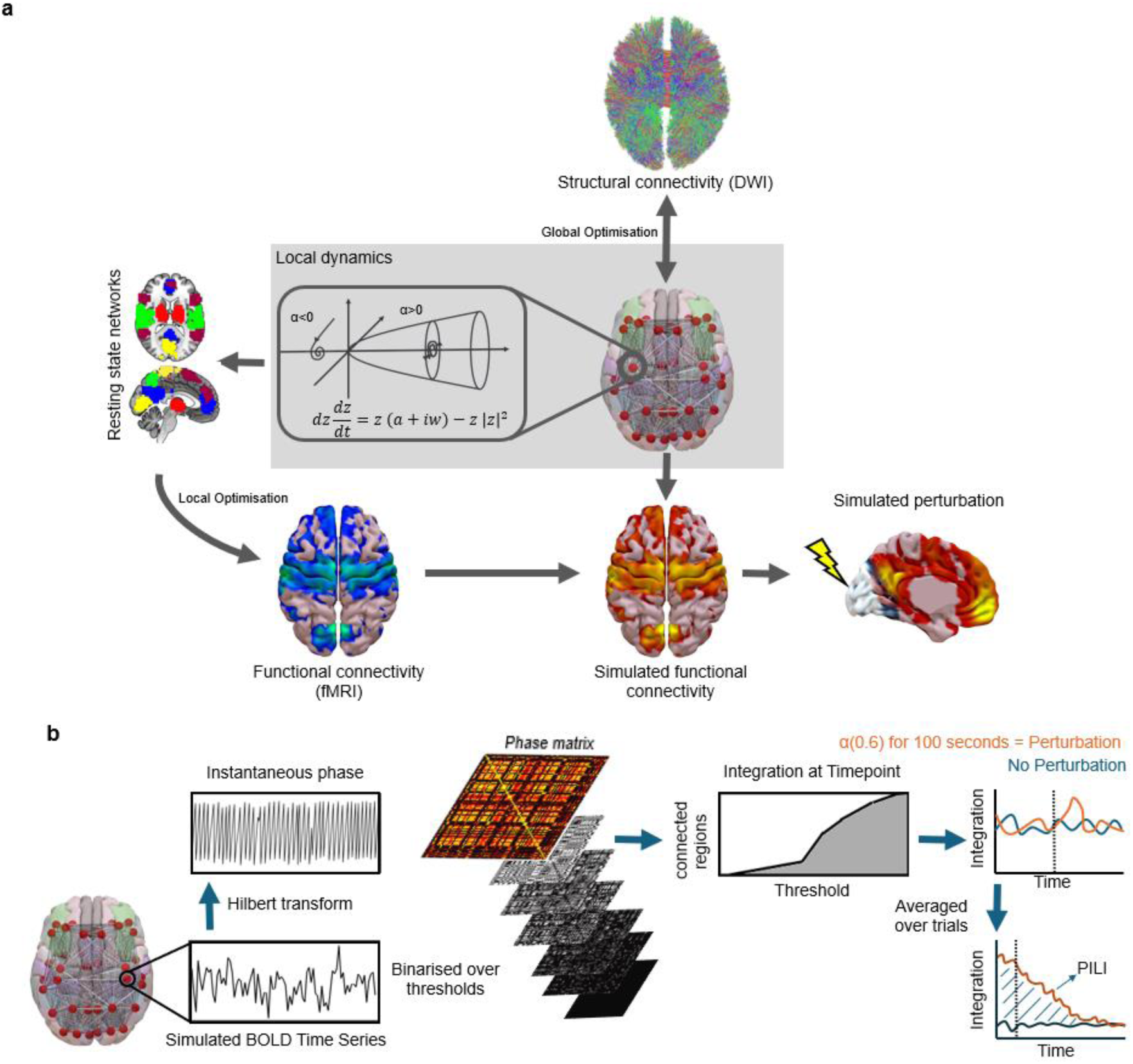
Schematic describing the main computational methods used. **a.** Construction of the whole-brain models. This model uses empirical structural and functional connectivity to simulate the BOLD time series. The local dynamics are defined by a Hopf bifurcation, which depending on the value of the bifurcation parameter can be at stable fixed-point (a<0), stable limit cycle (a>0) or a bifurcation between both regimes (a=0). The structural connectivity provided by white matter tractography from DWI modulates the coupling between brain regions and the global coupling parameter scales the coupling between brain regions in the SC. The local bifurcation parameters are optimised towards the empirical fMRI data, restricted to a parameter space representing 9 resting state networks obtained from an independent component analysis. Brain dynamics are assessed by simulating a perturbation and observing the resultant return to a dynamical baseline. **b.** PILI protocol. We applied the Hilbert transform to the simulated BOLD signals to obtain the instantaneous phases. We constructed and binarised a phase locking matrix at each time point and calculated the number of regions in the largest connected component over thresholds. The integration was defined as the integral over all thresholds. The perturbation protocol consisted of modifying the bifurcation parameter of one brain region to the stable regime (α=0.6) for 100 s. Integration was then calculated over 300 s in the basal unperturbed state and after immediately the model perturbation. PILI is computed as the integral between the curves of integration values over time for the perturbed dynamics (orange) compared to the maximum of the basal state dynamics (blue).

We then sought to investigate the dynamical properties of the different states of consciousness following an *in-silico* perturbation. Each model was exposed to a perturbation protocol, which consisted of shifting the dynamics of one of the 90 brain regions to a more stable state for 100s, repeated for each brain region. The integration over time was then determined immediately after the perturbation, and in a baseline state without perturbation. We then calculated PILI for each model after perturbation by summing the area beneath the perturbational integration curve until it returned to the baseline state (Figure 1b). We determined the PILI for each node by selectively perturbing one node while maintaining the dynamics constant for the other nodes across 100 trials. These trials were then averaged to produce a single PILI value for each node. Finally, a singular, mean PILI value representing the average across all brain regions was computed.

To statistically compare the PILI values between states of consciousness, we took each simulation as a separate datapoint, thus having 90 brain regions and 100 simulations for each region. We calculated the Cohen’s d effect size to compare the distributions between each condition and its associated comparison. In states of reduced consciousness, DoC patients and healthy subjects under anaesthesia had decreased PILI values compared to the comparison conditions (CNT and wakefulness, for DoC and anaesthesia groups respectively) (Figure 2). Specifically, UWS (Cohen’s d *=* -0.52) and MCS (Cohen’s d = -0.48) patients had lower PILI values compared with CNT. Similarly, decreases in PILI values were also observed for propofol (Cohen’s d = -0.54) and dexmedetomidine anaesthesia (Cohen’s d = -0.32). In contrast, we found that PILI values in the psychedelic conditions were higher compared to the placebo condition, for both LSD (Cohen’s d = 0.11) and psilocybin (Cohen’s d *=* 0.28). Network-wise changes in PILI are listed in Supplementary Table 2.

**Figure 2:**
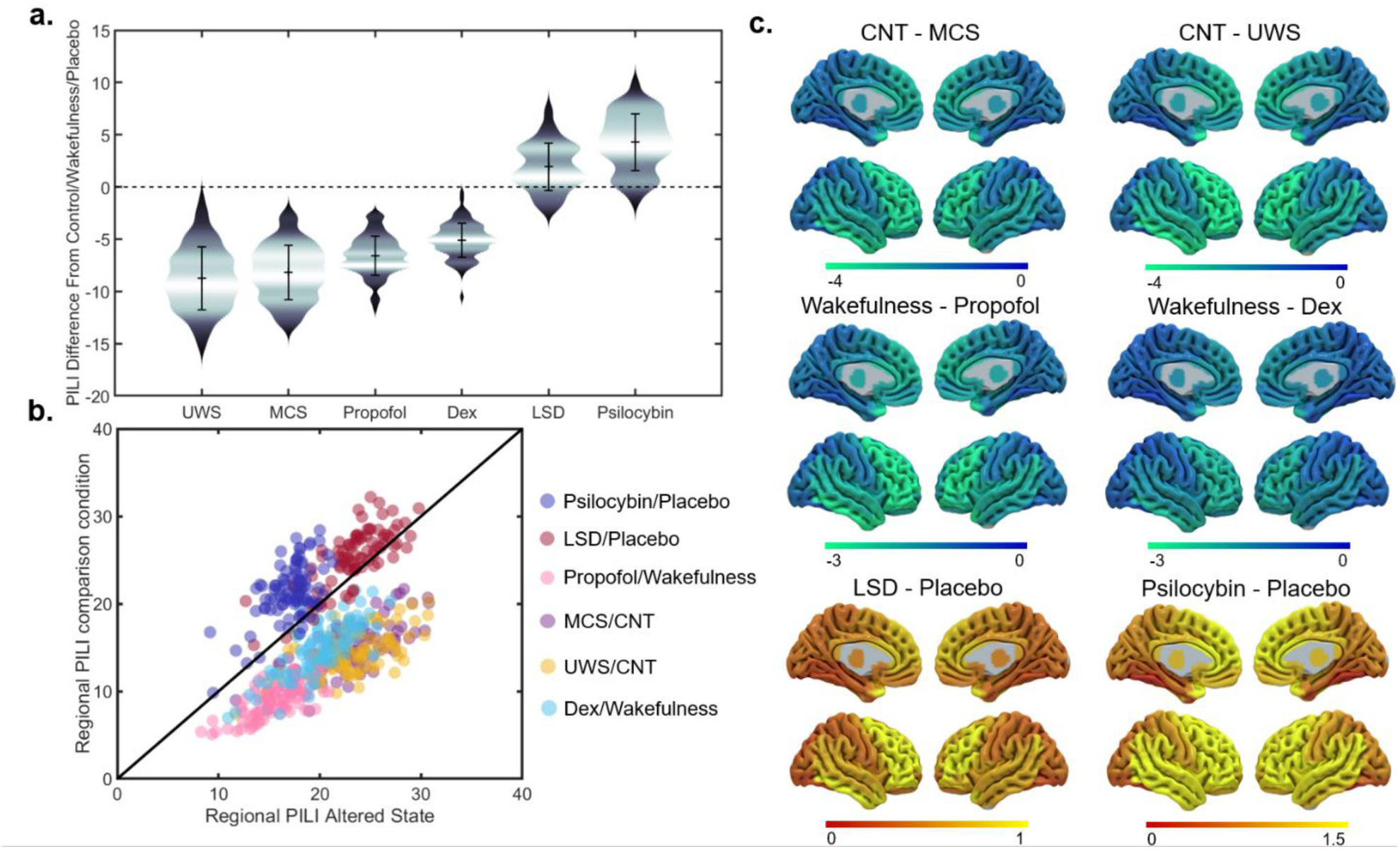
Brain dynamics assessed by PILI in whole-brain models of in different states of at the group level. **a.** Violin plots of the distribution of PILI trials averaged over all brain regions of group level models in each state of consciousness. Y axis represents the distance of each group from its respective comparison group: healthy controls for the UWS and MCS groups, the respective wakefulness conditions for propofol and dexmedetomidine, respective placebo conditions for LSD and psilocybin. **b.** Absolute region-wise PILI in each state of consciousness on the X axis plotted against the respective comparison group on the Y axis. **c.** Absolute network-wise changes in PILI.

### Perturbations of individualised patient models

We then constructed individualised whole-brain models for each subject in our DoC dataset (20 UWS, 26 MCS, 35 CNTs). The parameters of each model were optimised towards the individual BOLD fMRI data. To define the coupling between nodes, we used the individual patient structural connectivity (SC), generated by white matter tractography from the DWI data. From this point onwards, all the analyses were performed using these digital twins of each subject.

We performed the perturbation protocol on each patient individually and calculated PILI values for each of the 90 brain regions. In all our analyses on individual models, we decided to consider a single PILI value in each brain region as the mean across all 100 simulations. PILI values were larger in the CNT group compared to the UWS group (z=3.60, *p*<0.001). Additionally, PILI values were larger in the MCS group compared to the UWS group (z=2.34, *p*=0.02) (Figure 3).

**Figure 3:**
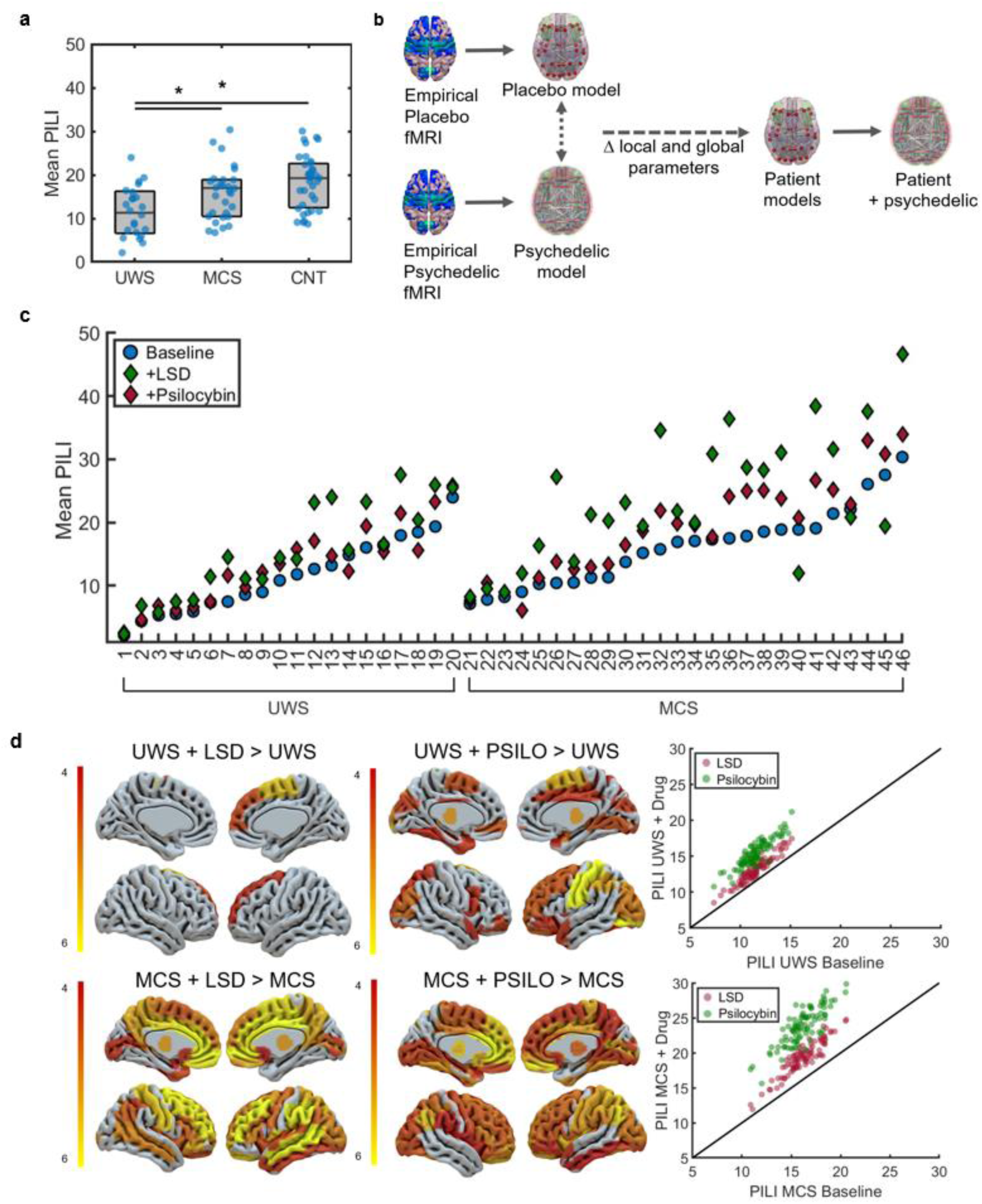
Brain dynamics assessed by PILI in personalised whole-brain models of individual patients with DoC before and after the simulation of LSD and psilocybin. **a.** Mean PILI in individual patient models and healthy controls. Each blue dot represents a patient or control. **b.** The virtual pharmacology method. Computational models of each state of consciousness in the same subjects are constructed. The global coupling and local bifurcation parameters are extracted and applied to the models built on individual patients to simulate the administration of LSD or psilocybin **c.** Individual patient models before and after simulation of LSD and psilocybin. Blue circles represent the mean PILI of each patient at baseline, red diamonds represent LSD and green diamonds represent psilocybin. **d.** Left: Plotted t-statistic values of region wise t-tests displaying brain regions with significant increases, in PILI from simulating LSD and psilocybin in patients with UWS (top) and MCS (bottom), Bonferroni corrected for multiple comparisons for the 90 brain regions. Right: Average PILI values for each of the 90 brain regions, averaged across patients at the baseline state (X-axis) and after the simulation of LSD (red) and psilocybin (green) (y-axis).

### Simulation of LSD and psilocybin on patients with DoC: A virtual clinical trial

We then simulated the administration of LSD and psilocybin on the individual patient models. This was achieved through firstly creating whole-brain models optimised at the group level using the BOLD fMRI data from healthy individuals being administered a drug (LSD, psilocybin) and at the respective placebo conditions. Within both the LSD and the psilocybin models individually, we extracted the changes in the global coupling and local bifurcation parameters that manifested upon comparing the placebo model to the psychedelic model. We then applied the shift in parameters representing the virtual administration of LSD and psilocybin separately to each patient’s model. The resulting psychedelic-induced changes in brain dynamics were then characterised by calculating PILI at the baseline state and after the simulation of the psychedelic drug. Please consult the Supplementary Figure 1 for a longer discussion about the validity of our virtual pharmacology approach.

Subsequently, we simulated the administration of LSD and psilocybin to each individual patient model. Simulating LSD significantly increased the PILI values of patients with DoC, with a greater effect on patients in the MCS (z=4.13, *p*<0.001) compared to patients in the UWS (z=2.8, *p*<0.001). This was also the case after simulating psilocybin, as the increases in PILI values observed in patients in the UWS (z=3.71, *p*<0.001) were lower than those observed in patients in the MCS (z=3.92, p<0.001). Next, we examined how the global response to perturbation differed across initial stimulation site, calculating the PILI values in each region, averaged across subjects at the baseline, and after simulating LSD and psilocybin. All regional differences in PILI values can be found in Supplementary Table 3. To better contextualise the regional changes in PILI, we calculated the differences in PILI in the 9 resting state networks, which define our local parameter space (Table 1). In the MCS group after the simulation of both LSD and psilocybin, the networks with the largest differences in PILI values were the DMN, the attention and limbic network. In the UWS group, the sensorimotor network had the highest increase in PILI values for both the LSD and psilocybin conditions. For both LSD and psilocybin, region-wise PILI values did not correlate with regional SC after Bonferroni correction for the 4 comparisons (LSD: MCS r=0.21, *p*=0.04, UWS r=0.05 *p*=0.56; psilocybin: MCS r=0.13, *p*=0.23; UWS r=0.02, *p*=0.84), nor FC (LSD: MCS r=0.17, *p*=0.11, UWS r=0.12, *p*=0.84; psilocybin: MCS *p*=0.26, r=0.02, UWS r=0.05, *p*=0.61).

**Table 1.** Network wise changes in PILI in each group and condition. Wilcoxon signed-rank tests. Bold indicates significance after Bonferroni correction for the 9 networks.

| Group |  | Brain networks |  |  |  |  |  |  |  |  |
| --- | --- | --- | --- | --- | --- | --- | --- | --- | --- | --- |
|  |  | Attention | Auditory | Default mode | Frontoparietal | Limbic | Precuneus | Sensorimotor | Thalamus | Visual |
| UWS+ LSD | Z-statistic | 2.77 | 2.69 | 2.61 | 2.23 | 2.69 | 2.46 | **2.95** | 2.73 | 2.73 |
|  | P-value | 0.005 | 0.007 | 0.008 | 0.028 | 0.007 | 0.014 | **0.003** | 0.006 | 0.006 |
| UWS+ Psilocybin | Z-statistic | **3.92** | **3.92** | **3.85** | **3.62** | **3.88** | **3.32** | **3.92** | **3.88** | **3.88** |
|  | P-value | **<0.001** | **<0.001** | **<0.001** | **<0.001** | **<0.001** | **<0.001** | **<0.001** | **<0.001** | **<0.001** |
| MCS + LSD | Z-statistic | **4.153** | **4.026** | **4.229** | **3.518** | **4.127** | **4.127** | **4.076** | **4.026** | **3.899** |
|  | P-value | **<0.001** | **<0.001** | **<0.001** | **<0.001** | **<0.001** | **<0.001** | **<0.001** | **<0.001** | **<0.001** |
| MCS + psilocybin | Z-statistic | **3.797** | **3.584** | **3.847** | **3.70** | **3.746** | **3.797** | **3.721** | **3.797** | **3.721** |
|  | P-value | **<0.001** | **<0.001** | **<0.001** | **<0.001** | **<0.001** | **<0.001** | **<0.001** | **<0.001** | **<0.001** |

### Baseline predictors of treatment response in individual patients

We then sought to identify which structural and functional capacities of patients’ brains at baseline were associated with a high simulated treatment effect. In this way, we could ascertain the most promising candidates for potential psychedelic treatment. We defined simulated treatment effect to be the difference in PILI between the baseline state and after the simulation of LSD and psilocybin. We quantified the strength of FC across the nine resting state networks that define our local bifurcation parameter space. Additionally, we computed several graph theory metrics from the patients’ SC matrices, including global efficiency, local efficiency, centrality, and graph strength. We also determined the mean fractional anisotropy (FA) values from DWI data. See Supplementary Table 4-5 for statistical analyses of the structural and functional capacities at baseline.

Here, we present 4 representative example patients and their structural and functional characteristics at baseline to elucidate insights into which characteristics could predict treatment efficacy (Figure 4a). These include one patient from each diagnostic group who had the large increases and decreases in PILI, averaged between psilocybin and LSD. MCS S41 showed the largest increase in PILI values for both LSD and psilocybin. At baseline, MCS S41 had one of the highest mean FC strengths of the group (z=4.46, *p<*0.001). However, their mean SC strength was not significantly different from the group mean (z=0.4, *p*=0.74). UWS S07 had one of the highest simulated treatment effects of the group. This patient had a mean FC which was not significantly different from the group mean (z=0.4291, *p*=0.70). However, the mean SC was significantly higher than the group mean (z=3.16, *p*=0.002). Examining those subjects with the worst simulated treatment outcome, UWS S14 had a relatively high FC (z=2.65, p=0.008), yet one of the worst SC of the group (z=-4.20, *p*<0.001). Whilst MCS S40 had a SC not significantly different from the mean of the group (z=0.79, *p*=0.43), yet a low mean FC (z=-2.98, *p*=0.002).

**Figure 4:**
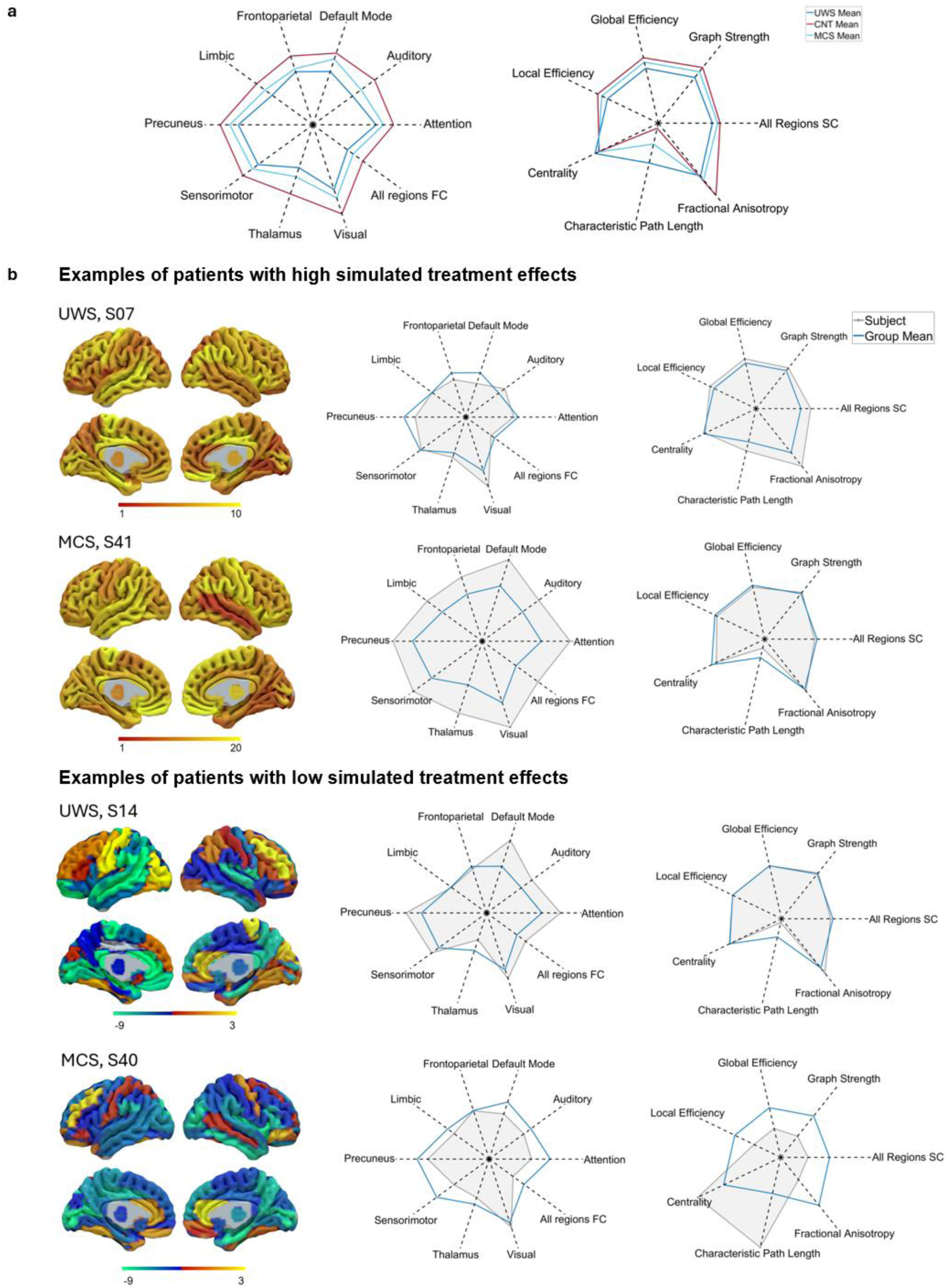
Structural and functional connectivity characteristics at baseline in the patients with DoC. **a.** Group average patient and healthy control graph theory structural metrics and functional connectivity in resting state networks at baseline**. b.** Examples of patients and their functional and structural capacities at baseline. One example is chosen from each diagnostic group that underwent the high and low average simulated treatment effect between psilocybin and LSD conditions. Left: Absolute regional changes in PILI values for each subject. Warmer colours (red and yellow) indicate higher PILI changes, while cooler colours (green and blue) indicate lower PILI changes. Right: Radar plots comparing the functional and structural capacities of each patient (filled grey shape) to the mean of the associated diagnostic group (blue line). The left radar plot illustrates the average FC across all regions and within the networks that define the local parameter space: frontoparietal, default mode, auditory, attention, visual, thalamus, sensorimotor, precuneus, and limbic networks. The right radar plot shows average SC metrics across all regions, and the average global efficiency, graph strength, local efficiency, betweenness centrality, fractional anisotropy, and characteristic path length.

To further investigate these relationships and identify baseline biomarkers of predicted treatment efficacy, we performed correlation analyses between each structural and functional capacity and the simulated treatment effect. In the UWS group, only the mean SC (LSD: r=0.67, *p*=0.001; psilocybin: r=0.65, *p*<0.001) correlated with the simulated treatment effect and remained significant after Bonferroni corrections (Figure 5). In the MCS group, there were no significant correlations between structural capacities and the simulated treatment effect for either drug (Supplementary Table 6). However, the mean FC strongly correlated with the simulated treatment effect for both LSD (r=0.65, *p*<0.001) and psilocybin (r=0.56, *p*=0.002) (Figure 5). Additionally, in both the LSD and psilocybin conditions, FC in the DMN significantly correlated with the simulated treatment effect, and for LSD, FC in the attention, limbic network, and sensorimotor network also correlated with the simulated treatment effect (Table 2).

**Figure 5:**
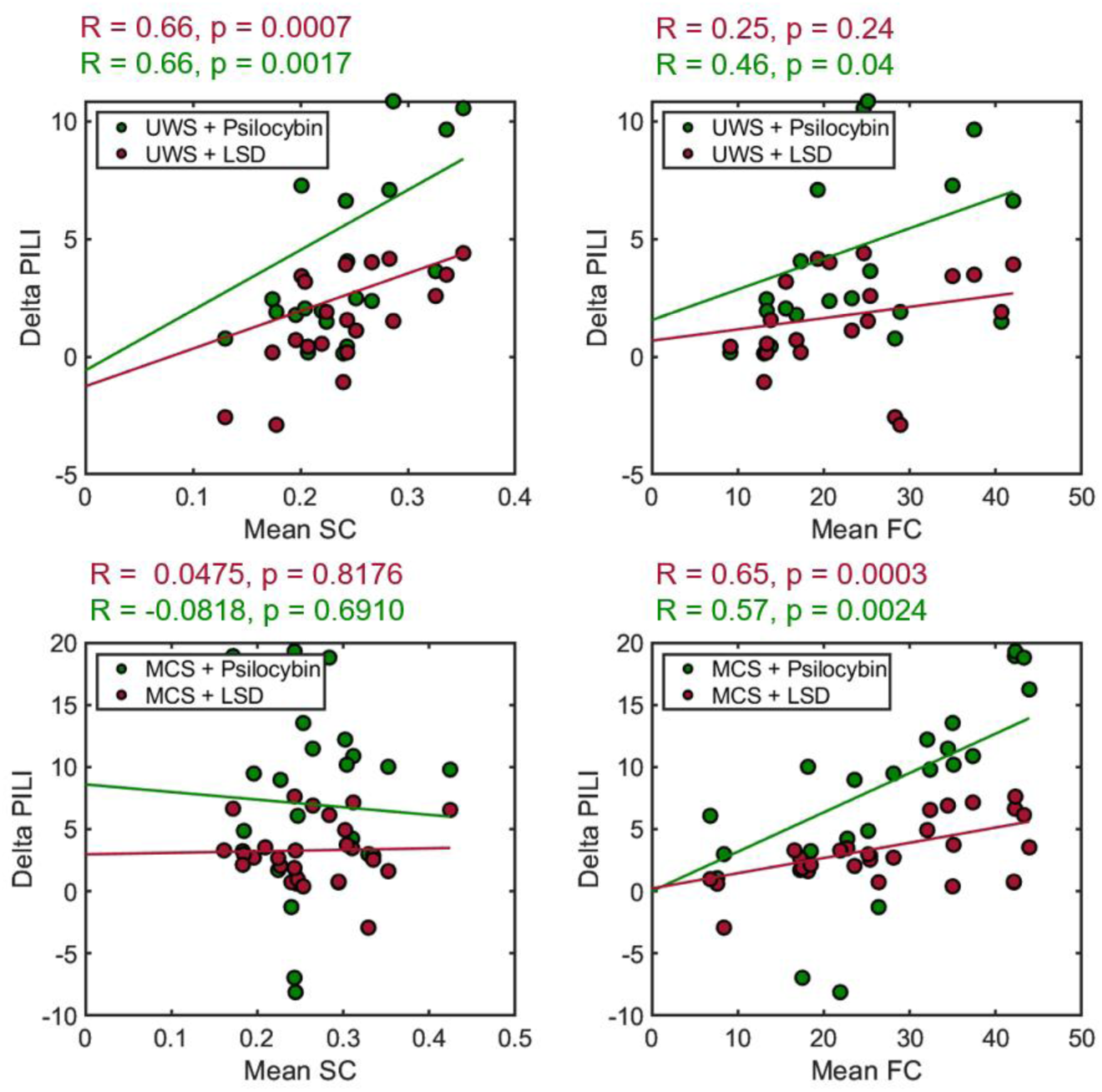
Baseline SC correlated with treatment response in patients with UWS, whereas baseline FC correlated with simulated treatment response in patients in the MCS. Scatter plots showing correlations between the changing in PILI values as a result of simulating LSD and psilocybin (Delta PILI) and the average baseline SC and FC in UWS patients (Left) and MCS patients (Right). Regression lines are indicated in red (LSD) and green (psilocybin). Associated correlation coefficients (R) and p-values are stated above in red (LSD) and green (psilocybin). Star indicates significance after Bonferroni correction for the 8 correlations.

**Table 2:** Correlation coefficients and p-values of correlational analyses between the strength of FC in each resting state network at baseline and the change in PILI as a result of simulating psilocybin and LSD in patients with UWS and MCS. Star represents significance after Bonferroni correction for the 9 networks.

| Group | Condition | Attention | Auditory | Default Mode | Frontoparietal | Limbic | Precuneus | Sensorimotor | Thalamus | Visual |
| --- | --- | --- | --- | --- | --- | --- | --- | --- | --- | --- |
| UWS + LSD | R-value | 0.035 | 0.184 | 0.225 | 0.199 | 0.340 | 0.159 | 0.050 | 0.353 | 0.395 |
|  | P-value | 0.885 | 0.438 | 0.334 | 0.401 | 0.142 | 0.502 | 0.834 | 0.127 | 0.085 |
| UWS + psilocybin | R-value | 0.203 | 0.348 | 0.365 | 0.371 | 0.465 | 0.152 | 0.140 | 0.328 | 0.451 |
|  | P-value | 0.391 | 0.132 | 0.114 | 0.107 | 0.034 | 0.522 | 0.558 | 0.158 | 0.046 |
| MCS + LSD | R-value | **0.622\*** | 0.466 | **0.567\*** | 0.298 | **0.569\*** | 0.521 | **0.592\*** | 0.531 | 0.266 |
|  | P-value | **0.0007\*** | 0.017 | **0.003\*** | 0.139 | **0.002\*** | 0.006 | **0.001\*** | 0.005 | 0.189 |
| MCS + psilocybin | R-value | 0.467 | 0.435 | **0.598\*** | 0.347 | 0.534 | 0.468 | 0.427 | 0.520 | 0.302 |
|  | P-value | 0.016 | 0.027 | **0.001\*** | 0.082 | 0.005 | 0.016 | 0.028 | 0.007 | 0.134 |

## Discussion

We used individualised whole-brain computational models in patients with DoC based on empirical DWI and fMRI data. Using each digital twin, we conducted a virtual clinical trial to explore the novel treatment paradigm of using psychedelic drugs, specifically LSD and psilocybin, to improve the brain dynamics of patients with DoC. To this end, we firstly set out to validate PILI, a modelling-based perturbational biomarker based on the integration of a modelled perturbation in distinguishing between states of consciousness namely, DoC, anaesthesia and the psychedelic state. Secondly, we simulated the administration of LSD and psilocybin on each patient and assessed the resulting brain dynamics as a proxy for simulated treatment response. Thirdly, we characterised the baseline predictive biomarkers associated with increases in PILI, ultimately identifying predictive biomarkers of potential treatment efficacy.

Assessing the brain’s response to a perturbation allows us to understand how it integrates external information and provides insights into to the criticality of brain dynamics and their implied complexity. Our results showed that PILI could distinguish between states of consciousness according to the phenomenological richness of experience. States of diminished consciousness such as DoC (UWS and MCS) and anaesthesia (propofol and dexmedetomidine) showed lower PILI values compared to the control subjects and wakefulness condition respectively. States with a higher richness of experience, i.e. the psychedelic state (LSD and psilocybin), possessed higher PILI values compared to their placebo conditions. This supports research linking brain complexity and criticality to consciousness^53^, ideas which are at the core of integrated information theory of consciousness^54,55^. States in which consciousness is diminished such as in DoC or anaesthesia, possess more stable brain dynamics, with lower complexity^15,16,19,20,35,56^. This dynamic rigidity, manifesting through a faster return to the baseline following a perturbation, exemplifies the shift away from criticality. Support from TMS-EEG studies on anaesthesia and DoC show lower evoked complexity in these states compared to normal wakefulness^43^. During the psychedelic state, the increase in PILI epitomises the brain’s shift even closer to criticality. This echoes several previous lines of investigation showing the psychedelic state is associated with an increase in brain complexity, an increased repertoire of available states and more critical dynamics^30–35^. Both propofol and dexmedetomidine were administered at unresponsive doses. However, dexmedetomidine often induces dreaming, which occurs less often with propofol^57^. This likely contributes to dexmedetomidine’s intermediate PILI value and underscores the importance of considering phenomenological differences between unresponsive states^58^.

When comparing PILI values from diagnostic groups at the single-subject level, there was a difference in mean PILI values between the UWS and MCS groups, but not between the MCS and CNT groups. This mirrors findings from studies on the PCI, where below a numerical cut-off, a patient is considered unconscious, and above this level there is less discrimination between conscious states like MCS and CNT^43,59^. It also aligns with clinical assessments of consciousness at the bedside where UWS patients do not show any signs of consciousness, yet MCS patients exhibit signs of residual consciousness. The established threshold allows PCI to function as a diagnostic tool at the single-subject level. While PILI currently lacks this discriminative capacity at the single subject level, future developments in computational modelling should aim to create a diagnostic tool of comparable precision.

We also developed a method to administer virtual pharmacological treatments based on extracting the changes in parameters observed due to the drug effects and applying them to models that represent other states of consciousness, here, patients with DoC. In principle, this approach could be generalised to extract and apply parameter changes that represent any pharmacological and neuromodulation intervention that acutely results in changes in brain connectivity, such as anaesthetics, stimulation techniques, or conditions e.g. listening to music. Crucially, the success of this method depends on the parameter extraction originating from a dataset which estimates each state within the same subjects. This ensures the most accurate estimation of the treatment effects by imposing the specificity of functional and structural changes. To illustrate the importance of this, we present (Supplementary Figure 2) and discuss (Supplementary Discussion 2) analyses using a separate fMRI dataset from healthy subjects under psilocybin, acquired with a between-subjects design. Here, the PILI was slightly lower in the psilocybin condition compared to the placebo condition and the simulated treatment response in patients in the UWS was not associated to the SC in patients with UWS.

The PILI increased in response to the administration of LSD and psilocybin to patients with DoC. This represents a shift in brain dynamics towards criticality, with greater effects observed in the MCS compared to UWS groups. The simulated psychedelic-induced changes in brain dynamics in MCS patients show a similar absolute shift towards criticality as those seen between UWS and CNT, and between anaesthesia and wakefulness. Whilst at face value, this may imply that the absolute size of the change in complexity could be enough to support a conscious state, the phenomenological implications and behavioural consequences of these changes in PILI remain an open question.

Considering region-specific changes in PILI approaches, the cogent question of how these changes in simulated brain dynamics might manifest phenomenologically; how might the treatment be experienced in-vivo? In MCS patients, after both LSD and psilocybin, the largest network-wise increases in PILI were observed in the DMN, attention, and limbic networks. Notably, the DMN and limbic networks also showed the largest absolute increases in PILI in group-level models of LSD and psilocybin in healthy subjects, which is also corroborated by a 2019 paper modelling perturbations in models of LSD^30^. Decreases in within-network integrity, particularly of the DMN, is a commonly reported finding in fMRI psychedelic literature^50,51,60–62^. Decoupling between medial temporal lobe (part of the limbic network) and the DMN, along with altered connectivity between the DMN, dorsal attention and salience network has been associated with ego dissolution^60,62,63^. This phenomenon is characterised by a diminished experience of selfhood, often with a feeling of environmental connectedness^64^ and positive feelings (e.g., joy, ecstasy, awe, peace)^65^. Curiously, patients with DoC, a population with a questionable level of selfhood, also exhibit disrupted DMN connectivity, with MCS patients having stronger DMN connectivity compared to UWS patients^66,67^. If similar psychedelic-induced alterations to functional network characterisation occurs in patients with DoC, as our results suggest, it may further impair the integrity of these networks. An interesting predictive speculation could be an increase of environmental sensitivity and perceptual processing driven by the global shift towards criticality, yet perhaps an accompanied by a reduction of residual selfhood^37^, and the ability to outwardly focus attention, driven by increased integration between the DMN, attention, and limbic networks. Vitally, any increase in environmental awareness could be life-changing for patients and caregivers and it remains difficult to predict how exactly the interaction between environmental sensitivity, selfhood and cognitive capacity may materialise. In fact, in MCS patients, the DMN, limbic, and attention networks showed the highest baseline connectivity correlation to the simulated treatment effect. A phenomenological consequence of this may imply a strong self-model and attention resources underwritten by high DMN, attention, and limbic network FC, are necessary for any treatment response. In the UWS group, the only network with significant changes in PILI for both psilocybin and LSD was the SM network. This is likely due to the large cortical structural damage present in other networks, restricting the propagation of activity into such areas. The restriction of increases in PILI to the SM network, decreases the likelihood of treatment outcomes positively augmenting consciousness and cognition in patients in the UWS.

Whilst generally consistent, psilocybin and LSD had some differential effects, with psilocybin showing more variability and a greater effect in UWS patients. Although both drugs produce similar phenomenological effects^68^, they have distinct pharmacological profiles: LSD targets D1-3 receptors, while psilocin, the active metabolite of psilocybin, inhibits the serotonin transporter^69^. Psilocybin and LSD have been shown to have unique patterns of functional reorganisation dependent on distinct neurotransmitter mechanisms^70^. Also, recent modelling work simulating the stimulation of different receptors in patients with DoC has demonstrated the differential effects these receptors have on brain dynamics. Compared to 5-HT2A receptor stimulation, activation of D1 and D2 receptors had minimal impact on the modelled trajectory of brain dynamics in patients with DoC towards that of controls^71^. These differences in pharmacology and connectivity likely contribute to the variations in their simulated effects on brain dynamics and highlight their potential differing utility in treatments.

In UWS patients, the SC drove the simulated treatment response, whereas FC was shown to be more pertinent in MCS patients. Neither the network connectivity nor average FC correlated with simulated treatment response in UWS patients, whilst in MCS patients none of the structural characteristics correlated with treatment response. This suggests that MCS patients may already have a sufficiently connected structure, which liberates the SC from limiting the predicted treatment response. Since patients in the MCS have more connecting white matter fibres which support a richer FC at rest, the psychedelic-induced increases in complexity could be facilitated through utilising less commonly engaged pathways and networks in the brain that are nonetheless structurally present. In contrast, UWS patients lack the structural basis from which the psychedelic-induced increases in complexity could resonate. This outlines a hierarchical relationship between structure, function and complexity. With biological structure being the foundational dependency upon which functional dynamics build; complexity cannot emerge from an absence of structure. The mean SC was more strongly associated to treatment outcome rather than any specific graph measure. This suggests that the severity of structural damage present in patients in the UWS means that only an estimate of sheer number of white matter fibres dictates the preserved capacity to support complex brain dynamics, rather than any other network characteristic. These results represent a grim outlook for UWS patients with large structural damage. However, there is a growing interest in another use case of psychedelics in those with severe brain injury that seeks to harness the neuroplastic effects to augment recovery immediately in the acute phase of brain injury^72^. Future work utilising models that account for plasticity could explore this. Taken together, our correlational analyses suggest that an MCS patient with high average FC and specifically within the DMN and attention networks has the greatest likelihood to have the largest changes in brain dynamics following a psychedelic drug.

This study has several limitations. The inherent range in PILI values within each subject at the single-subject level underscores the limited applicability of PILI for between-subject comparisons, emphasising its more appropriate use for characterising states within the same subjects. Similarly, differences in acquisition protocols and scanning parameters make comparing PILI values between datasets challenging, as evidenced by the variability in PILI values across different control group models (see Supplementary Figure 3 and Supplementary discussion 2). The Hopf model is a phenomenological model that does not aim to replicate underlying biology. However, recent studies have shown that many complex bottom-up models at the regional level merely replicate a Hopf bifurcation^73^. Therefore, whilst the Hopf model is limited in its mechanistic explanatory power compared to biophysical models, they still similarly capture regional and global brain dynamics. In addition, this work relies on the superposition assumption, being that a linear subtraction of the effects of psychedelics on controls well predicts the action of the drugs in patients with DoC. Despite this significant assumption, it provides a useful approximation in the absence of empirical data. Additionally, our method is based off parameter estimations from a single dataset which may introduce bias.

Here, we ran a virtual clinical trial by simulating the administration of LSD and psilocybin in individualised whole-brain models of patients with DoC. We showed that psychedelics increase the brain’s response to perturbation and globally, bring the brain dynamics of patients with DoC closer to criticality. We also revealed that patients in the MCS are more likely to increase in complexity compared to UWS patients and that this is dependent upon the strength of the baseline FC. This work builds on idea of using psychedelics for DoC and contributed towards computational personalised medicine and drug discovery.

## Methods

### Dataset descriptions

This study includes fMRI data from 6 datasets, 3 from the University and University Hospital of Liège (DoC, propofol, dexmedetomidine), 2 from Imperial College London (LSD, psilocybin) and 1 from Maastricht University (psilocybin). For the DoC dataset, we also used DWI acquired in the same patients and healthy controls.

#### DoC dataset

The study received approval from the Ethics Committee of the Faculty of Medicine at the University of Liège. Written informed consent was obtained from the healthy control participants and from the legal representatives of the patients. We included 46 adult patients with DoC, comprising 26 in MCS (6 females, age range 23–74 years; mean age ± SD, 47 ± 17 years), and 20 in UWS (6 females, age range 31–74 years; mean age ± SD, 52 ± 16 years), along with 35 gender-matched healthy controls (14 females, age range 19–72 years; mean age ± SD, 40 ± 14 years). Data for the DoC patients were recorded 57.98 ± 203.65 months post-injury. There are 10 acute patients with DoC (4 UWS, 5 MCS) in the sample. The diagnosis for DoC patients was confirmed through repeated behavioural assessments using the Coma Recovery Scale-Revised (CRS-R) conducted by trained staff, which evaluates auditory, visual, motor, sensorimotor function, communication, and arousal^74^. DoC patients were included if their MRI exams were conducted without anaesthetic sedation and if they had undergone at least five CRS-R assessments within a 14-day period with one assessment conducted the same day as the MRI acquisition. Patients were excluded based on the following criteria: (i) having any significant neurological, neurosurgical, or psychiatric disorders prior to the brain injury leading to DoC, (ii) having contraindications to MRI such as implanted electronic devices or external ventricular drains, and (iii) being medically unstable or having extensive focal brain damage, defined as damage affecting more than two-thirds of one hemisphere. Further details on the demographics and clinical characteristics of the patients are provided in Supplementary Table 1. Structural and functional MRI (fMRI) data were acquired on a Siemens 3T Trio scanner (Siemens Inc, Munich, Germany). The BOLD fMRI resting state (i.e. task free) was acquired using EPI, gradient echo with following parameters: volumes = 300, TR = 2000 ms, TE = 30 ms, flip angle = 78∘, voxel size = 3 × 3 × 3 mm3, FOV = 192 × 192 mm2, 32 transversal slices, with a duration of 10 minutes. Subsequently, structural 3D T1-weighted MP-RAGE images were acquired with following parameters: 120 transversal slices, TR = 2300 ms, voxel size = 1.0 × 1.0 × 1.2 mm3, flip angle = 9∘, FOV = 256 × 256 mm2). The diffusion MRI (dMRI) data were collected using a single echo planar imaging sequence gradient scheme with 64 non-collinear gradient directions (total acquisition time [TA] = 14:47 min, b = 1,000 s/mm², field of view [FOV] = 256 mm², voxel size = 2.0 mm isotropic, repetition time [TR] = 9,700 ms, echo time [TE] = 92 ms).

#### Propofol and dexmedetomidine datasets

The propofol and dexmedetomidine datasets have been previously published^48,49^. Both datasets have identical experimental protocols and received approval from the Ethics Committee of the Medical School of the University of Liege (University Hospital, Liege, Belgium). After receiving informed consent from the volunteers, sixteen healthy control subjects (age range 18-31 years) underwent propofol-induced sedation. Eleven healthy control subjects (age range 19-29 years) underwent dexmedetomidine-induced sedation. fMRI data were collected during normal wakefulness with eyes closed and during propofol/dexmedetomidine-induced sedation. Propofol/dexmedetomidine was infused through an intravenous catheter placed in a vein of the right hand or forearm, and an arterial catheter was placed in the left radial artery. Sedation was achieved using a computer-controlled intravenous infusion of propofol to maintain constant effect-site concentrations (for details on the procedure, see^48^). The drug plasma and effect-site concentrations were estimated using a three-compartment pharmacokinetic model. After reaching the appropriate effect-site concentration, a 5-minute equilibration period was allowed to ensure equilibration of drug distribution between compartments. Arterial blood samples were taken immediately before and after the scan in each clinical state for subsequent determination of the drug concentration. The level of consciousness was clinically evaluated throughout the study using the Ramsay scale^75^. Responsiveness was assessed through volitional hand squeezing in response to a verbal command from the experimenter. When there was no response to the command, the subject was considered to be in deep sedation (Ramsay 5–6). For each assessment of consciousness level, the Ramsay scale verbal commands were repeated twice. Before and after each scanning session, a reaction time task was also performed to provide additional information on the clinical state of the volunteers. The healthy subjects had no MRI contraindications, no history of neurological or psychiatric disorders, and no drug consumption that could significantly affect brain function. The propofol and dexmedetomidine datasets were acquired on a 3T Siemens Allegra scanner (Siemens AG, Munich, Germany). The fMRI resting-state scans were acquired using the following parameters: EPI, gradient echo, volumes = 200; TR = 2460 ms, TE = 40 ms, voxel size = 3.45 × 3.45 × 3 mm³, FOV = 220 × 220 mm, 32 transverse slices, 64 × 64 × 32 matrix size. The structural images were acquired using 3D T1-weighted MP-RAGE with the following parameters: 120 transverse slices, TR = 2250 ms, TE = 2.99 ms, voxel size = 1 mm³, flip angle = 9°, FOV = 256 × 240 × 160 mm.

#### LSD dataset

We used previously published data fully described here^50^. Briefly, 12 healthy participants were scanned under six different conditions: LSD resting state, placebo resting state, LSD and placebo resting state while listening to music, and LSD and placebo resting state after listening to music. LSD and placebo sessions were separated by at least 14 days, with the condition order balanced across participants, who were blinded to this order. For this study, we only used the LSD resting state and placebo resting state data. All participants provided informed consent. The experimental protocol was approved by the UK National Health Service Research Ethics Committee, West-London. The experiments conformed to the revised Declaration of Helsinki, the International Committee on Harmonisation Good Clinical Practice guidelines, and the National Health Service Research Governance Framework. Each participant received either 75 µg of LSD (intravenous, I.V.) or saline/placebo (I.V.) 70 minutes prior to MRI scanning. Participants reported noticing subjective drug effects between 5- and 15-minutes post-dosing. The drug effects peaked between 60- and 90-minutes post-dosing. The subsequent plateau of drug effects varied among individuals, generally lasting for four hours post-dosing. MRI acquisition started approximately 70 minutes post-dosing and lasted about 60 minutes. After each of the three scans, participants performed subjective ratings inside the scanner via a response box. The scans conducted with saline/placebo were considered baseline MRI scans compared to the LSD scans. Neuroimaging data were collected using a 3T GE HDx MRI system. Data were recorded using a gradient echo-planar imaging sequence, TR/TE = 2000/35 ms, field of view = 220 mm, 64 × 64 acquisition matrix, parallel acceleration factor = 2, 90° flip angle.

#### Psilocybin Imperial dataset

We used previously published data described here^51^. Briefly, 15 healthy volunteers participated in two MRI scanning sessions separated by at least 14 days. Participants were at least 21 years old, with no personal or family history of major psychiatric disorders, no substance dependence, no cardiovascular disease, and no history of adverse responses to psychedelic drugs. All subjects had prior experience with psilocybin but had not used it within 6 weeks of the study. The study received approval from a National Health Service research ethics committee, and informed consent was obtained from all participants. During each session, participants received either psilocybin (2 mg dissolved in 10 mL saline, administered via a 60-second intravenous injection) or a placebo (10 mL saline, administered via a 60-second intravenous injection) in a counterbalanced design. Infusions commenced precisely 6 minutes after the start of the 12-minute fMRI scans. Psilocybin’s effects were immediate and persisted throughout the scanning session. Neuroimaging data were collected using a 3T GE HDx MRI system. Anatomical scans preceded functional scans and were thus performed before drug or placebo administration. Structural scans were acquired using 3D fast spoiled gradient echo sequences in an axial orientation with a field of view of 256 × 256 × 192 and a matrix of 256 × 256 × 192, yielding 1 mm isotropic voxel resolution (repetition time/echo time TR/TE = 7.9/3.0 ms; inversion time = 450 ms; flip angle = 20). BOLD-weighted fMRI data were acquired at 3T using a gradient echo EPI sequence with TR/TE = 3000/35 ms, a field of view of 192 mm, a 64 × 64 acquisition matrix, a parallel acceleration factor of 2, and a 90° flip angle. Fifty-three oblique axial slices were acquired in an interleaved fashion, each 3 mm thick with no slice gap (3 × 3 × 3-mm voxels). Following the same motion exclusion criteria as the LSD dataset, nine subjects (seven men, mean age 32 ± 8.9 years) were included in the analysis.

#### Psilocybin Maastricht dataset

We analysed previously published data, described in detail here^52^. Briefly, data from 49 healthy volunteers with previous experience with a psychedelic drug not within the 3 months prior to the study. Following a randomised, placebo-controlled, double-blind parallel group design, participants were assigned to one of two conditions (0.17 mg/kg psilocybin, or placebo) such that groups were matched for age, sex, and educational level. The drug was administered orally in a closed cup containing bitter lemon (placebo) or bitter lemon and psilocybin (powder). Informed consent was obtained from all subjects.The study obtained ethical approval from the Maastricht University’s Medical Ethics Committee and was in accordance with the Medical Research Involving Human Subjects Act (WMO) as well as with the code of ethics on human experimentation from the declaration of Helsinki. Following criteria outlined in the original publication, one subject was excluded due to having a maximum framewise displacement of >0.75mm (half-voxel size). Four subjects were excluded on the basis of <5 minutes of the scan remaining after scrubbing. One further subject was excluded due to missing BOLD values in several brain regions. The final sample was 22 in the psilocybin group and 26 subjects in the placebo group. Participants underwent structural MRI 50-min post psilocybin/placebo administration, in addition to 6-min resting-state fMRI 102-min post psilocybin/placebo administration during the peak subjective drug effects. All images were acquired in a MAGNETOM 7T MRI scanner using the following acquisition parameters: TR = 1,400 ms; TE = 21 ms; field of view = 198 mm; flip angle = 60°; oblique acquisition orientation; interleaved slice acquisition; 72 slices; slice thickness = 1.5 mm; voxel size = 1.5 mm, isotropic. Furthermore, participants were presented with a black fixation cross on a white background during the scanning session and were asked to focus on the cross, clear their minds, and lie as still as possible.

### Functional magnetic resonance imaging and functional connectivity estimation

Since the datasets were acquired at different locations, we endeavoured to mitigate scanner and acquisition differences by re-preprocessing all data with the same pipeline and following uniform denoising procedures. The preprocessing of fMRI images was conducted utilising FSL (FMRIB Software Library v6.0, Analysis Group, FMRIB, Oxford, UK). Structural T1-weighted images were processed using the fsl_anat function, which involves brain extraction, bias field correction, and normalisation. Functional MRI data underwent preprocessing with MELODIC (Multivariate Exploratory Linear Optimized Decomposition into Independent Components). The Maastricht psilocybin fMRI data uniquely underwent realignment and susceptibility distortion correction with FSL_topup to correct for magnetic field distortion caused by the high strength 7T field. Main preprocessing steps for all datasets included discarding the initial 5 volumes, motion correction with MCFLIRT, brain extraction using BET (Brain Extraction Tool), spatial smoothing with a 5 mm FWHM Gaussian kernel, rigid body registration, and applying a high-pass filter with a cutoff of 100.0 seconds. Additionally, single-session ICA with automatic dimensionality estimation was performed in order to identify noise driven components. Each component was visually inspected using the FSLeyes package in Melodic mode to classify the single-subject Independent Components as either signal or noise/lesion-driven artifacts, based on the spatial map, time series, and temporal power spectrum. Noise components were then regressed out using fsl_regfilt to obtain a cleaned version of the functional data. Furthermore, BOLD data was detrended, demeaned, and band-pass filtered between 0.01 to 0.08 Hz. To obtain the BOLD time series for the 90 cortical and subcortical brain regions defined by the AAL atlas (excluding the cerebellum), FSL tools were used in each individual’s native EPI space. This parcellation has been found to be particularly suitable for studying whole-brain spatiotemporal dynamics^30,44,76^. Given the computational models were personalized at the individual level, utilising 90 regions provided a balanced approach between spatial resolution and computational feasibility. The cleaned functional data were registered to the T1-weighted structural image using FLIRT. The T1-weighted image was then normalised to the standard MNI space using FLIRT (12 DOF) and FNIRT. The resulting transformations were concatenated, inverted, and applied to warp the resting-state atlas from MNI space to the cleaned functional data. Nearest-neighbour interpolation ensured the preservation of labels. The BOLD time series for each of the 90 brain regions were extracted for each subject in their native space using fslmaths to create a binary mask of each brain region and fslmeants to obtain the time series of each mask. Each of the 90 regional average BOLD signals were then filtered in the 0.04-0.07 Hz range, which has previously been used due to its functional relevance and resistance to noise^77–80^. Pairwise Pearson correlation coefficients were then computed between all 90 brain regions for each subject. For the group level matrices, the Fisher transform was applied to the r-values to derive z-values for the 90×90 FC matrices before being averaged and subsequently back transformed.

### Anatomical connectivity

The patient structural connectomes were created using advanced, single-subject analyses of the diffusion MRI (dMRI) dataset of patients with DoC including 35 age-matched controls. MRI scans were performed using a Siemens 3T Trio scanner (Siemens Inc, Munich, Germany) with a 64-channel head coil. Using tools from MRtrix3 (https://.mrtrix.org/), raw dMRI data were corrected^81,82^ denoised and corrected for Gibbs ringing artefacts^83^, as well as distortions induced by motion, eddy currents, and EPI/susceptibility artefacts^84,85^. The response functions from the pre-processed diffusion-weighted images were estimated using the Dhollander algorithm and used to calculate fibre orientation distributions (FODs) via constrained spherical deconvolution (CSD)^86^. Here, we utilised MRtrix3Tissue (v5.2.8; https://3tissue.github.io), a variant of MRtrix3^87^, which enables 3-tissue constrained spherical deconvolution, generating separate FODs for white matter (WM), grey matter (GM), and cerebrospinal fluid (CSF) SS3T-CSD^88^, from single-shell (+b=0) diffusion MRI data. Whole-brain tractography was then performed^89^, generating 20 million streamlines per subject^90^ using dynamic seeding. The spherically informed filtering of tractograms (SIFT2) algorithm was applied to align the fibre density of the reconstructed streamlines with the underlying white matter structures^89,91^. Compared to tractograms constructed solely by the number of streamlines, SIFT2 adjusts the weight of individual streamlines to ensure alignment with the underlying image data^89,91,92,92^. SIFT2 is highly reproducible^93^ and enhances the biological interpretability of the estimated white matter tracts^94^. We then performed probabilistic tractography using the iFOD2 (improved 2nd order integration over fibre orientation distributions) algorithm, which enhances anatomical plausibility^95^. During tracking, the direction for each step is obtained by sampling from the FOD at the current position, with the probability of a particular direction being proportional to the FOD amplitude. Additional tractography settings included a default step size of 0.8 mm, a maximum angle between successive steps of 45°, a maximum length of 250 mm, a minimum length of 5 mm, a cutoff FA value of 0.08, and the use of b-vectors and b-values from the diffusion-weighted gradient scheme in FSL format. Concurrently, the b0 image in native space was registered to the T1 anatomical image using FSL FLIRT^96,97^. The T1 structural image was then normalised to standard space using FLIRT and FNIRT. The resulting transformations were reversed and applied to the AAL atlas in standard MNI space using a nearest neighbour algorithm to convert the atlas to each subject’s native space. The number of streamlines connecting each pair of regions was then estimated to produce the final structural connectivity matrix for each patient. To summarise, for each participant, a 90 x 90 symmetric weighted network of structural connectivity (SC) was constructed which represents the density of white matter tracts between anatomical regions of the brain. The fractional anisotropy (FA) values were generated by firstly estimating the diffusion kurtosis tensor map from the pre-processed DWI image. A map of tensor derived FA for each patient was then averaged across voxels to produce one FA value per subject. Graph theory metrics were calculated using The Brain Connectivity Toolbox^98^ implemented in MATLAB (version R2017a, MathWorks, Natick, MA).

We calculated graph strength, global efficiency, characteristic path length, average local efficiency, and betweenness centrality to capture different aspects of network structure. These measures provide insights into node importance, communication efficiency, and overall network topology.

### Computational whole-brain model

We used a whole-brain model to simulate the brain activity of each of the 90 cortical and subcortical brain regions in the AAL atlas. In particular, the dynamics of each region were described by means of coupled Stuart-Landau oscillators in the normal form of supercritical Hopf bifurcations^30,42,44,45^. This model simulates spontaneous signals akin to the average BOLD signal in a given brain region. The coupling between oscillators representing each region is driven by a given structural connectivity. The model at node *j* is described by the following set of coupled differential equations:

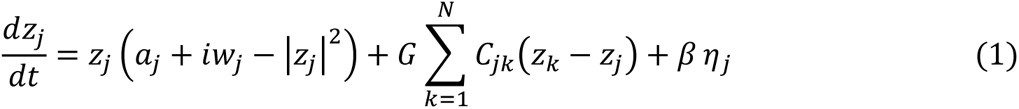

where *z* is a complex variable *z*^*j*^ = *x*_*j*_ + *iy*_*j*_, *w*_*j*_ is the natural node frequency (estimated as the average peak frequency of each region of the BOLD recordings of each group or subject, for the group and single-subject models respectively), *η*_*j*_ is additive Gaussian noise, *β* is the s scaling factor for the Gaussian noise (set to 0.04), *C* is the SC matrix, where *C*_*jk*_ is the coupling between nodes *j* and *k*, *G* is the global coupling that acts as a scaling factor for the SC matrix for all nodes, and *α*_*j*_ is the bifurcation parameter controlling the behaviour of the oscillator. For *α*∼0 the system is at a supercritical Hopf bifurcation and the Gaussian noise induces complex dynamics as the system oscillates between both sides of the bifurcation. For *α* < 0 the system decays to a stable fixed point at *z*_*j*_ = 0 and noise-induce oscillations appear due to the Gaussian noise. For *α* > 0 the system dynamics set into a stable limit cycle with a frequency of 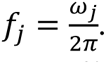 The SC matrix was scaled to a maximum value of 0.2 to ensure oscillatory dynamics bifurcations^30,42,44,45^. The fMRI BOLD signal simulation was obtained as the real part of *z* (*x*_*j*_). The stochastic differential equations were solved by means of the Euler-Maruyama method.

For all our group models involving healthy subjects, including those for psychedelics, anaesthesia, and the associated wakefulness and placebo comparisons, we used the SC matrix representing the average of the healthy controls in the DoC group. This approach was necessary as we did not have access to the individual SC data for each subject in each dataset. In previous modelling research a template SC obtained from a commonly used dataset is usually employed which represents the white matter connectivity of healthy subjects. This is assumed to well represent the SC of healthy controls in general^30,41,42,42,44,45,99^. Here, we adopt the same approach by using the SC estimate from the healthy control group of the DoC dataset. This ensured that the strength of the connectivity values, and thus the optimal parameters for the models employing a healthy structure, were in the same range as those of patients with DoC.

Crucially, our models of patients with DoC included, for the first time, the empirical structural data for each patient. The diverse aetiology of patients with DoC often results in significant structural differences between patients. By using personalised SC, we avoided the inherent bias of assuming a healthy white matter structure in DoC patients, enabling us to build more robust models incorporating more empirical information. For the first time, we created individualised single-subject whole-brain models using the individual structural and functional data.

### Model fitting to empirical data

This work involved the creation of 7 models at the group level and 48 models at the single subject level. For the group level analyses, the FC averaged over each subject in each condition was used to obtain the parameters for each model. For the single-subject analyses the individual FC of each patient was used to fit their parameters. Two parameters were fitted to the models in two steps. First, the global coupling *G* was fitted to the FC matrix by exploring a parameter range in steps of 0.01 starting from 0 until the fitting curve reached an optimum value. Three hundred seconds of BOLD activity were simulated each time. The same filtering applied to the empirical data was applied to the simulated time series and their FC matrices were obtained. Afterwards we optimised G based on the Kolmogorov-Smirnov distance (KS distance) between the empirical and simulated FC. For each *G* value, 500 simulations were performed and the curve of the average KS distance was used the determine the optimal *G* value at the local minimum.

Using the KS distance, the distribution of the FC values of the simulations were brought to a similar level to that of the empirical data. After fixing the *G* values for each group/subject as a starting point, in a second step the *α* values were also fitted in order to fine tune their dynamics by adjusting the bifurcation parameters of the regions. For each region *j* the bifurcation parameter *α*_*j*_ was obtained as the linear combination of 9 resting state networks including that region, obtained by means of an independent component analysis (ICA) of the fMRI data from the healthy controls in the DoC dataset. The ICA networks identified were the default mode, auditory, attention, visual, sensorimotor, precuneus, limbic, frontoparietal and thalamus. Thus, the parameter space was reduced to a dimension of 9. We used the Euclidean distance as the goodness of fit parameter to optimise the fit for each region, and a genetic algorithm to find the optimal values of the *α* parameters of each network as in ^42,100^. We ran the algorithm 10 times and used the average *α* values of the networks over the course of the 10 runs as the final optimal value. Ten was chosen as a balance between computational power requirements and finding the true optimal value. For each network, the parameter range of its associated *α* values to explore by the genetic algorithm was set to -0.2 to 0.2 ^42^.

### Integration

Integration refers to the capacity of the brain to assimilate communication between discrete processing units. The integration measure used in this study evaluates the synchronisation of brain activity measured by the BOLD signal and has been previously defined in ^30,44^. The Hilbert transform was applied to filtered BOLD signals to extract the instantaneous phase *φ*_*j*_(*t*) for each region j. The Hilbert transform derives the analytic representation of a real-valued signal, in this case, the BOLD time series. The synchronisation between pairs of brain regions was characterised as the difference between their instantaneous phases. At each time point, the phase difference P_jk_(t) between two regions j and k was calculated as

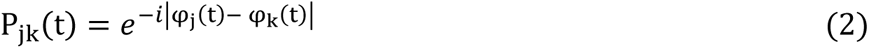

Here, *P*_*jk*_ (*t*) = 1 when the two regions are in phase, φ_j_(t) = φ_k_(t) (full synchronised). At any time *t*, the phase interaction matrix *P(t)* represents the instantaneous phase synchrony of each pair of brain regions. The phase interaction matrix is then binarised over 100 evenly spaced thresholds between 0 and 1. Then, the size of the largest connected component across all brain regions is obtained at each threshold, thus creating a function of connected components as a function of the threshold. The integration at time *t* is then denoted as the integral of this curve (Figure 1b).

### Model perturbation protocol and Perturbative Integrative Latency Index

The perturbation protocol has been previously described in^30,101^. Briefly, we used the local bifurcation parameters of the Hopf model to simulate a perturbation through switching each α from its original base value to a more synchronous regime (*α* = 0.6) for 100 s. We computed the integration over 400 s of simulated BOLD time series in the basal state and immediately after the perturbation offset. This protocol was performed at every brain region in the 90 AAL parcellation and repeated 100 times. After perturbation, the bifurcation parameter was reset to its original value obtained from the model fitting procedure. PILI in a particular region was calculated as the area under the integration curve from perturbation offset to the interception point with the basal state integration. This is repeated for each brain region for 100 trials before being averaged to obtain a regional PILI value. The singular mean PILI, represents the average PILI over all brain regions. Since the PILI is based on the integral of the integration curve, it characterises both the strength and latency of the return to its basal state after a model perturbation.

### Virtual pharmacology approach

To simulate the administration of psilocybin and LSD to the individualised patient models, we developed a virtual pharmacology method. This was based upon group level models optimised to the BOLD fMRI data from healthy individuals being administered a drug (LSD, psilocybin) and at the respective placebo conditions. The global coupling parameters and local bifurcation parameters representing each condition were then extracted and compared within each condition to attain a shift in the local bifurcation parameters which represented that particular condition. This change in parameters thus represented an instructional change in how to shift the model dynamics from one condition to another. We then applied this shift in parameters representing the virtual administration of the drug (LSD and psilocybin) separately to each patient’s model.

### Statistical analyses

When comparing the mean PILI values between whole-brain models built at the group level, we used Cohen’s d effect size to compare the distributions of PILI values produced by each model. Since each group level model can in theory, produce an infinite number of simulations, thus artificially inflating the sample size, any p-values produced by frequentist statistical tests are less applicable in these cases.

For all analyses using individual models, we employed a standard frequentist approach since, each model essentially can act as a unique datapoint with its own distribution, akin to a subject. To compare the PILI values between diagnostic groups, from the personalised models of DoC patients, we performed Mann-Whitney *U* tests. To compare the mean PILI values obtained from simulating the psychedelic drug to the baseline PILI values, we performed Wilcoxen signed-rank tests. We used paired t-tests to evaluate the region-wise changes in PILI as a result of simulating the psychedelic with Bonferroni correction for the 90 regions. Wilcoxon signed-rank tests were used to compare the network-wise PILI values between the baseline state and simulated drug state with Bonferroni correction for each of the 9 resting state networks.

## Conflict of interest

At the time of publication, VB has a financial relationship with Orion Pharma, Edwards Medical, Medtronic, Grünenthal, and Elsevier.

## Acknowledgements

This work was supported by the University and University Hospital of Liège (Liège, Belgium), the Belgian National Fund for Scientific Research (FRS-FNRS), the ERA-Net FLAG-ERA JTC2021 project ModelDXConsciousness (Human Brain Project Partnering Project), FLAG-ERA JTC2023 project BrainAct, the Fondation Léon Fredericq, the FNRS MIS project (F.4521.23), the FNRS PDR project (T.0134.21), the GIGA Doctoral School for Health Sciences, the Mind Science Foundation, the fund Generet, the King Baudouin Foundation, the BIAL Foundation and the DoC-Box project (HORIZON-MSCA-2022-SE-01-01-101131344). PC and NA are research fellows, OG is research associate, and SL is research director at the F.R.S.-FNRS. JA is postdoctoral fellow at the FWO (1265522N). We thank the patients and their families and the control subjects for participating in our study.

## Notes

### Competing Interest Statement

VB has a financial relationship with Orion Pharma, Edwards Medical, Medtronic, Grunenthal, and Elsevier.

