## Supplementary Materials for "A virtual clinical trial of psychedelics to treat patients with disorders of consciousness"

#### Demographic Information

| Subjects | Aetiology | TSI (days) | Age | Gender | Best CRS-R total score |
|---|---|---|---|---|---|
| UWS 1 | NTBI | 29 | 44 | M | 4 |
| UWS 2 | NTBI | 50 | 69 | F | 5 |
| UWS 3 | TBI | 8 | 48 | M | 2 |
| UWS 4 | TBI | 283 | 52 | M | 6 |
| UWS 5 | NTBI | 92 | 74 | M | 4 |
| UWS 6 | NTBI | 2890 | 49 | M | 6 |
| UWS 7 | TBI | 24 | 58 | M | 4 |
| UWS 8 | NTBI | 38 | 50 | F | 3 |
| UWS 9 | NTBI | 434 | 52 | M | 4 |
| UWS 10 | NTBI | 129 | 49 | F | 4 |
| UWS 11 | TBI | 3406 | 31 | M | 6 |
| UWS 12 | NTBI | 18 | 20 | M | 3 |
| UWS 13 | NTBI | 304 | 60 | M | 6 |
| UWS 14 | NTBI | 335 | 40 | F | 6 |
| UWS 15 | NTBI | 43 | 64 | M | 5 |
| UWS 16 | TBI | 2013 | 31 | M | 4 |
| UWS 17 | NTBI | 30 | 44 | M | 5 |
| UWS 18 | NTBI | 26 | 82 | M | 4 |
| UWS 19 | NTBI | 40 | 74 | F | 7 |
| UWS 20 | NTBI | 1683 | 39 | F | 7 |
| MCS 1 | NTBI | 389 | 59 | F | 8 |
| MCS 2 | Mixed | 35 | 73 | M | 5 |
| MCS 3 | TBI | 3034 | 34 | F | 12 |
| MCS 4 | NTBI | 20 | 52 | M | 14 |
| MCS 5 | TBI | 319 | 73 | M | 9 |
| MCS 6 | TBI | 21 | 25 | M | 15 |
| MCS 7 | TBI | 28 | 65 | M | 13 |
| MCS 8 | NTBI | 13 | 62 | M | 7 |
| MCS 9 | TBI | 135 | 51 | M | 12 |
| MCS 10 | TBI | 521 | 28 | M | 10 |
| MCS 11 | TBI | 1294 | 40 | M | 11 |
| MCS 12 | TBI | 25 | 67 | M | 11 |
| MCS 13 | TBI | 641 | 23 | F | 10 |
| MCS 14 | NTBI | 64 | 29 | M | 7 |
| MCS 15 | NTBI | 1482 | 32 | M | 14 |

| | | | | | |
|---|---|---|---|---|---|
| MCS 16 | TBI | 407 | 31 | M | 9 |
| MCS 17 | NTBI | 2639 | 38 | M | 9 |
| MCS 18 | TBI | 246 | 30 | M | 9 |
| MCS 19 | NTBI | 242 | 46 | F | 7 |
| MCS 20 | NTBI | 18 | 74 | F | 5 |
| MCS 21 | TBI | 134 | 66 | M | 16 |
| MCS 22 | TBI | 1331 | 35 | M | 8 |
| MCS 23 | TBI | 2690 | 24 | M | 14 |
| MCS 24 | NTBI | 9900 | 39 | M | 15 |
| MCS 25 | TBI | 674 | 66 | F | 7 |
| MCS 26 | NTBI | 396 | 57 | M | 8 |

**Supplementary Table 1:** Demographic information of MCS and UWS patients. The table includes condition, aetiology (traumatic brain injury (TBI), non-traumatic brain injury (NTBI)), time since injury (TSI), age, gender (F=female, M=male), best Coma Recovery Scale-Revised (CSR-R) total score.

#### PILI values from group level models

| Condition | UWS – CNT | MCS – CNT | Propofol – wakefulness | Dexmedetomidine – wakefulness | LSD – placebo | Psilocybin – Placebo |
|---|---|---|---|---|---|---|
| Whole brain Average | -0.52 | -0.48 | -0.54 | -0.32 | 0.11 | 0.28 |
| Attention | -2.37 | -2.07 | -2.26 | -1.29 | 0.41 | 1.36 |
| Auditory | -2.25 | -2.02 | -2.37 | -1.36 | 0.41 | 1.14 |
| Default Mode | -2.62 | -2.46 | -2.41 | -1.54 | 0.62 | 1.55 |
| Frontoparietal | -1.48 | -1.56 | -1.78 | -1.17 | 0.38 | 0.23 |
| Limbic | -3.5 | -2.57 | -3.23 | -2.03 | 0.78 | 2.01 |
| Precuneus | -1.25 | -1.2 | -1.18 | -0.69 | 0.4 | 0.57 |
| Sensorimotor | -1.89 | -1.65 | -1.77 | -1.08 | 0.42 | 1.11 |
| Thalamus | -2.84 | -2.44 | -2.81 | -1.81 | 0.43 | 1.18 |
| Visual | -2.04 | -2.14 | -1.93 | -1.32 | 0.29 | 0.74 |

**Supplementary Table 2:** Whole brain average and network-wise differences in PILI values from the group level models to the respective comparison condition, assessed by Cohen's d effect size.

#### Model validation

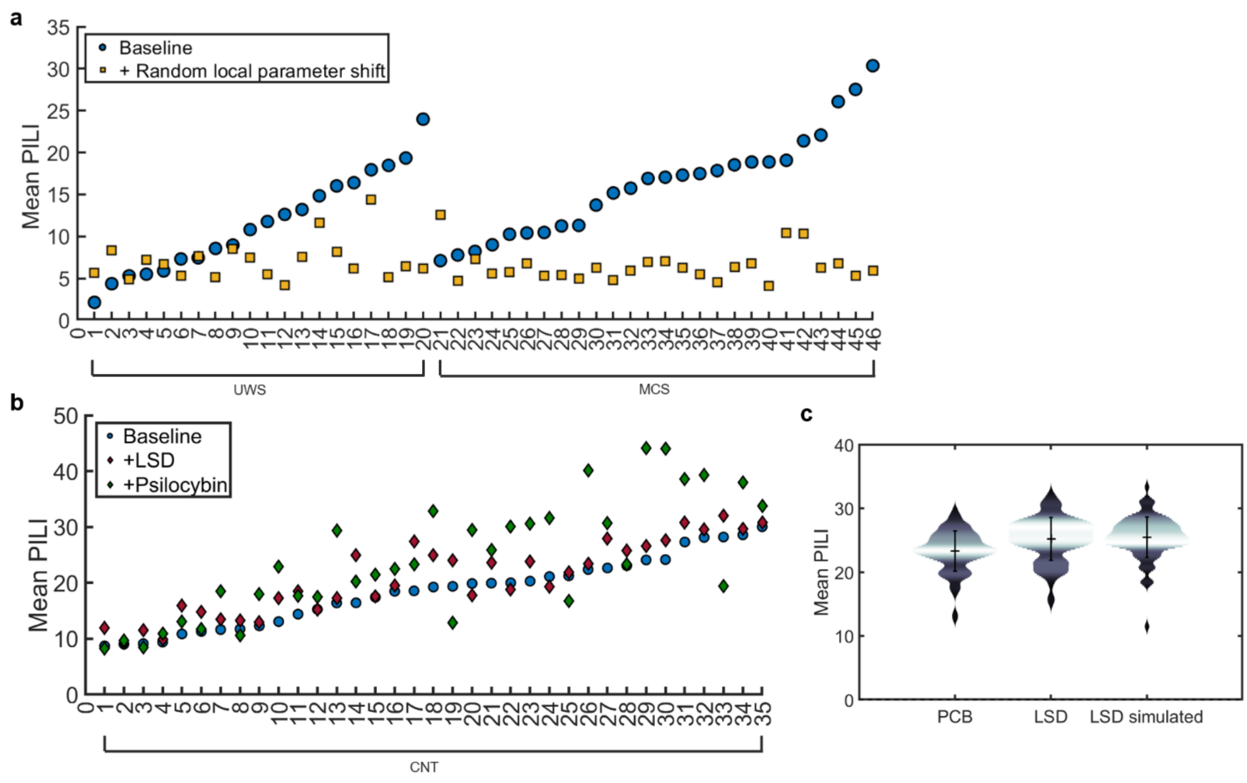

**Supplementary Figure 1: a**. Mean PILI values in single patient models individually perturbed by randomly shifting the alpha parameters between -0.3 and +0.3. **b.** Mean PILI values in individual models based on healthy controls. Red diamonds represent LSD and green diamonds represent psilocybin **c.** Mean PILI values from the group level models optimised to Placebo, LSD and after simulating the administration of LSD on the PCB model.

**Supplementary Discussion 1**: Model validation. To build evidence for our virtual pharmacology approach, we performed 2 additional perturbational analyses. Firstly, we demonstrated the importance of the distinct distribution of local parameters in dictating the dynamics of the model. Instead of applying the changes in bifurcation parameters, which represent the difference in between the drug state and the placebo state, each of the nine bifurcation parameters in each single patient model corresponding to each of the functional networks was modified by a random addition between -0.3 and +0.3. We then calculated PILI in each patient at baseline, and after a random shift in alpha parameters. In almost all cases, shifting the alpha parameters randomly, resulted in decreases in PILI values (Supplementary Figure 1a). This displays the vital importance of local bifurcation parameters in maintaining the complex dynamics of these models. We also simulated the administration of LSD and psilocybin on 35 single subject models optimised to our healthy control BOLD fMRI and DWI dataset (Supplementary Figure 1b). We showed that almost all subjects have increases in PILI from simulating LSD and psilocybin. Also, as with the group level models based on the empirical data, the simulation of psilocybin had a greater effect than LSD. Finally, we created a group-level model optimised to the BOLD fMRI placebo condition and then applied the changes in local and global parameters extracted from the LSD and PCB models (Supplementary Figure 1c). The rationale behind this was in an attempt to compare the PILI values obtained from the group level model with that obtained from simulating the LSD using our virtual pharmacology method. The simulation of LSD on the psilocybin model resulted in a mean PILI value of 25.47, similar to the modest increase in PILI of 25.21, obtained directly form the LSD model. This evidences the methods capacity to simulate the drug condition.

### Region-wise PILI values from individual level models

| UWS + psilocybin | | | UWS + LSD | | | MCS + psilocybin | | | MCS + LSD | | |
|---|---|---|---|---|---|---|---|---|---|---|---|
| **Brain region** | **Z-stat** | **p-value** | **Brain region** | **Z-stat** | **p-value** | **Brain region** | **Z-stat** | **p-value** | **Brain region** | **Z-stat** | **p-value** |
| ‘Fusiform_R’ | 2.087518038 | 0.047191265 | 'Lingual_R' | 0.526282381 | 0.60478175 | 'Fusiform_R' | 2.087518038 | 0.047191265 | 'Heschl_R' | 1.54986303 | 0.13374228 |
| 'Lingual_R' | 2.407343564 | 0.023777537 | 'Fusiform_L' | 0.620187566 | 0.542501994 | 'Lingual_R' | 2.407343564 | 0.023777537 | 'Amygdala_R' | 1.94254859 | 0.06341578 |
| 'ParaHippocampal_R' | 2.60147745 | 0.015373385 | 'Frontal_Inf_Tri_R' | 0.902932662 | 3.78E-01 | 'ParaHippocampal_R' | 2.60147745 | 0.015373385 | 'Fusiform_R' | 2.03978734 | 0.05207128 |
| 'Cingulum_Post_R' | 2.74445331 | 0.011056085 | 'SupraMarginal_L' | 0.936951804 | 0.360541268 | 'Cingulum_Post_R' | 2.74445331 | 0.011056085 | 'SupraMarginal_R' | 2.10968161 | 0.04506716 |
| 'Occipital_Mid_R' | 2.805121685 | 0.009593734 | 'Postcentral_L' | 1.007970898 | 0.326133646 | 'Occipital_Mid_R' | 2.805121685 | 0.009593734 | 'Lingual_R' | 2.164465 | 0.04017933 |
| 'Fusiform_L' | 3.100236295 | 0.004739812 | 'Occipital_Mid_L' | 1.097474992 | 0.286148352 | 'Fusiform_L' | 3.100236295 | 0.004739812 | 'Occipital_Inf_R' | 2.45392312 | 0.02144279 |
| 'Cuneus_R' | 3.113771868 | 0.00458649 | 'Cingulum_Post_R' | 1.144365848 | 0.266684414 | 'Cuneus_R' | 3.113771868 | 0.00458649 | 'Temporal_Inf_L' | 2.50761232 | 0.01901515 |
| 'Frontal_Inf_Oper_L' | 3.187776446 | 0.003828903 | 'Cingulum_Post_L' | 1.197115726 | 0.245984714 | 'Frontal_Inf_Oper_L' | 3.187776446 | 0.003828903 | 'Occipital_Mid_R' | 2.51130921 | 0.01885773 |
| 'Occipital_Sup_R' | 3.249469427 | 0.003290839 | 'Temporal_Mid_L' | 1.214323905 | 0.239501796 | 'Occipital_Sup_R' | 3.249469427 | 0.003290839 | 'ParaHippocampal_R' | 2.57480621 | 0.01633592 |
| 'Hippocampus_R' | 3.352989623 | 0.002547881 | 'Putamen_R' | 1.238020837 | 0.230788494 | 'Hippocampus_R' | 3.352989623 | 0.002547881 | 'Frontal_Mid_Orb_L' | 2.67170717 | 0.0130862 |
| 'ParaHippocampal_L' | 3.376894065 | 0.002400982 | 'ParaHippocampal_L' | 1.239366408 | 0.230301124 | 'ParaHippocampal_L' | 3.376894065 | 0.002400982 | 'Temporal_Mid_R' | 2.68825165 | 0.01259589 |
| 'Occipital_Inf_R' | 3.3916748 | 0.002314297 | 'Hippocampus_R' | 1.263868433 | 0.221564176 | 'Occipital_Inf_R' | 3.3916748 | 0.002314297 | 'Temporal_Pole_Mid_R' | 2.80694273 | 0.0095528 |
| 'SupraMarginal_R' | 3.521260239 | 0.001673766 | 'Temporal_Inf_R' | 1.289226376 | 0.212794487 | 'SupraMarginal_R' | 3.521260239 | 0.001673766 | 'Heschl_L' | 2.88480694 | 0.00794931 |
| 'Temporal_Pole_Mid_L' | 3.55683454 | 0.001530567 | 'Angular_L' | 1.295440176 | 0.210687358 | 'Temporal_Pole_Mid_L' | 3.55683454 | 0.001530567 | 'Temporal_Pole_Sup_R' | 2.93986517 | 0.00697342 |
| 'Thalamus_R' | 3.584233018 | 0.001428498 | 'Occipital_Inf_L' | 1.352522142 | 0.192087117 | 'Thalamus_R' | 3.584233018 | 0.001428498 | 'ParaHippocampal_L' | 3.1125774 | 0.00459983 |
| 'Cuneus_L' | 3.627040127 | 0.001282195 | 'Occipital_Sup_R' | 1.361607743 | 0.189250482 | 'Cuneus_L' | 3.627040127 | 0.001282195 | 'Rolandic_Oper_R' | 3.12442659 | 0.00446916 |
| 'Precentral_L' | 3.666010859 | 0.001161815 | 'Rolandic_Oper_R' | 1.371764703 | 0.186118952 | 'Precentral_L' | 3.666010859 | 0.001161815 | 'Temporal_Pole_Mid_L' | 3.15455122 | 0.00415282 |
| 'Occipital_Mid_L' | 3.686182468 | 0.001103925 | 'Olfactory_R' | 1.424225462 | 0.170598706 | 'Occipital_Mid_L' | 3.686182468 | 0.001103925 | 'Frontal_Mid_Orb_R' | 3.23940336 | 0.00337335 |
| 'Paracentral_Lobule_L' | 3.717712249 | 0.001019053 | 'Frontal_Sup_Orb_R' | 1.445073812 | 0.164728842 | 'Paracentral_Lobule_L' | 3.717712249 | 0.001019053 | 'Cingulum_Post_R' | 3.27525101 | 0.00308829 |
| 'Amygdala_R' | 3.818101712 | 0.000789276 | 'Hippocampus_L' | 1.450554231 | 0.163213385 | 'Amygdala_R' | 3.818101712 | 0.000789276 | 'Cuneus_R' | 3.32733621 | 0.00271521 |
| 'Cingulum_Post_L' | 3.877149836 | 0.000678783 | 'Amygdala_L' | 1.463469836 | 0.159686833 | 'Cingulum_Post_L' | 3.877149836 | 0.000678783 | 'Temporal_Sup_L' | 3.37152651 | 0.00243323 |
| 'Temporal_Inf_R' | 3.886806003 | 0.000662225 | 'Temporal_Sup_L' | 1.483981372 | 0.154214407 | 'Temporal_Inf_R' | 3.886806003 | 0.000662225 | 'Caudate_R' | 3.39039965 | 0.00232165 |
| 'Occipital_Sup_L' | 3.897536087 | 0.000644292 | 'Caudate_R' | 1.505018599 | 0.148762597 | 'Occipital_Sup_L' | 3.897536087 | 0.000644292 | 'Cingulum_Post_L' | 3.40600941 | 0.00223313 |
| 'Postcentral_L' | 3.935065601 | 0.000585252 | 'Rectus_L' | 1.514513463 | 0.146354563 | 'Postcentral_L' | 3.935065601 | 0.000585252 | 'Angular_L' | 3.40740727 | 0.00222537 |
| 'Lingual_L' | 4.022798118 | 0.000467258 | 'Fusiform_R' | 1.5348095 | 0.141315174 | 'Lingual_L' | 4.022798118 | 0.000467258 | 'Frontal_Inf_Orb_L' | 3.40829864 | 0.00222043 |
| 'Rolandic_Oper_R' | 4.037799059 | 0.000449583 | 'Angular_R' | 1.564892402 | 0.134111481 | 'Rolandic_Oper_R' | 4.037799059 | 0.000449583 | 'Hippocampus_R' | 3.46435703 | 0.00193038 |
| 'Caudate_R' | 4.052312286 | 0.000433112 | 'Putamen_L' | 1.642205571 | 0.116995442 | 'Caudate_R' | 4.052312286 | 0.000433112 | 'Hippocampus_L' | 3.48721658 | 0.00182299 |
| 'Insula_R' | 4.070384746 | 0.000413433 | 'Frontal_Med_Orb_R' | 1.693878988 | 0.10661869 | 'Insula_R' | 4.070384746 | 0.000413433 | 'SupraMarginal_L' | 3.520991 | 0.0016749 |
| 'Frontal_Inf_Orb_L' | 4.105012772 | 0.000378167 | 'Occipital_Inf_R' | 1.724647464 | 0.100821977 | 'Frontal_Inf_Orb_L' | 4.105012772 | 0.000378167 | 'Postcentral_L' | 3.52888381 | 0.00164202 |
| 'Olfactory_R' | 4.135105431 | 0.00034995 | 'Thalamus_L' | 1.746963324 | 0.096789257 | 'Olfactory_R' | 4.135105431 | 0.00034995 | 'Frontal_Inf_Oper_R' | 3.57050152 | 0.00147879 |
| 'Precuneus_R' | 4.148796688 | 0.000337812 | 'Frontal_Mid_Orb_L' | 1.787434055 | 0.089830634 | 'Precuneus_R' | 4.148796688 | 0.000337812 | 'Temporal_Sup_R' | 3.59250746 | 0.001399 |
| 'Occipital_Inf_L' | 4.210381557 | 0.000288175 | 'Amygdala_R' | 1.811056388 | 0.085973278 | 'Occipital_Inf_L' | 4.210381557 | 0.000288175 | 'Parietal_Inf_R' | 3.59677542 | 0.00138402 |

| | | | | | | | | | | | |
|---|---|---|---|---|---|---|---|---|---|---|---|
| 'SupraMarginal_L' | 4.240929857 | 0.00026631 | 'Cuneus_R' | 1.812425024 | 0.085754284 | 'SupraMarginal_L' | 4.240929857 | 0.00026631 | 'Temporal_Inf_R' | 3.69861365 | 0.00106967 |
| 'Supp_Motor_Area_R' | 4.241398147 | 0.000265988 | 'Occipital_Sup_L' | 1.838590297 | 0.081660304 | 'Supp_Motor_Area_R' | 4.241398147 | 0.000265988 | 'Occipital_Inf_L' | 3.72155749 | 0.00100915 |
| 'Temporal_Pole_Mid_R' | 4.273939798 | 0.000244533 | 'Frontal_Mid_R' | 1.840941752 | 0.081300902 | 'Temporal_Pole_Mid_R' | 4.273939798 | 0.000244533 | 'Precentral_R' | 3.82168768 | 0.00078209 |
| 'Temporal_Pole_Sup_L' | 4.293791307 | 0.000232298 | 'Temporal_Inf_L' | 1.872732743 | 0.076576349 | 'Temporal_Pole_Sup_L' | 4.293791307 | 0.000232298 | 'Pallidum_R' | 3.8332496 | 0.00075935 |
| 'Frontal_Mid_Orb_L' | 4.297149247 | 0.000230289 | 'Pallidum_L' | 1.8888787 | 0.074270694 | 'Frontal_Mid_Orb_L' | 4.297149247 | 0.000230289 | 'Angular_R' | 3.89136793 | 0.00065454 |
| 'Hippocampus_L' | 4.318576364 | 0.000217873 | 'Calcarine_L' | 1.987453006 | 0.061480348 | 'Hippocampus_L' | 4.318576364 | 0.000217873 | 'Cuneus_L' | 3.98234778 | 0.00051841 |
| 'Frontal_Sup_Medial_R' | 4.325541752 | 0.000213982 | 'Precuneus_R' | 1.993477884 | 0.060766467 | 'Frontal_Sup_Medial_R' | 4.325541752 | 0.000213982 | 'Frontal_Inf_Oper_L' | 3.99407665 | 0.00050304 |
| 'Calcarine_L' | 4.334443676 | 0.000209109 | 'Caudate_L' | 2.034082 | 0.05614586 | 'Calcarine_L' | 4.334443676 | 0.000209109 | 'Olfactory_R' | 4.00115181 | 0.00049398 |
| 'Rectus_L' | 4.337182864 | 0.000207632 | 'Postcentral_R' | 2.040439664 | 0.055451679 | 'Rectus_L' | 4.337182864 | 0.000207632 | 'Amygdala_L' | 4.04799271 | 0.00043795 |
| 'Postcentral_R' | 4.352760237 | 0.000199428 | 'Temporal_Mid_R' | 2.144771622 | 0.045113403 | 'Postcentral_R' | 4.352760237 | 0.000199428 | 'Calcarine_L' | 4.22891373 | 0.00027471 |
| 'Temporal_Mid_R' | 4.369655912 | 0.000190894 | 'Cingulum_Ant_R' | 2.146973903 | 0.044915414 | 'Temporal_Mid_R' | 4.369655912 | 0.000190894 | 'Parietal_Sup_R' | 4.23621232 | 0.00026958 |
| 'Temporal_Inf_L' | 4.374118849 | 0.000188701 | 'Paracentral_Lobule_R' | 2.166640638 | 0.043182125 | 'Temporal_Inf_L' | 4.374118849 | 0.000188701 | 'Olfactory_L' | 4.2567998 | 0.00025561 |
| 'Precentral_R' | 4.395183533 | 0.000178684 | 'Cuneus_L' | 2.238968252 | 0.037320299 | 'Precentral_R' | 4.395183533 | 0.000178684 | 'Lingual_L' | 4.28970002 | 0.00023477 |
| 'Cingulum_Ant_R' | 4.397245651 | 0.000177733 | 'Frontal_Sup_Medial_L' | 2.293914616 | 0.033365897 | 'Cingulum_Ant_R' | 4.397245651 | 0.000177733 | 'Temporal_Pole_Sup_L' | 4.31032655 | 0.00022257 |
| 'Temporal_Pole_Sup_R' | 4.430571502 | 0.000163035 | 'Temporal_Sup_R' | 2.303914344 | 0.032689229 | 'Temporal_Pole_Sup_R' | 4.430571502 | 0.000163035 | 'Fusiform_L' | 4.32405548 | 0.00021481 |
| 'Precuneus_L' | 4.488313776 | 0.000140378 | 'SupraMarginal_R' | 2.332955602 | 0.030795331 | 'Precuneus_L' | 4.488313776 | 0.000140378 | 'Occipital_Sup_L' | 4.36826506 | 0.00019158 |
| 'Supp_Motor_Area_L' | 4.488958908 | 0.000140144 | 'Paracentral_Lobule_L' | 2.338173801 | 0.030465999 | 'Supp_Motor_Area_L' | 4.488958908 | 0.000140144 | 'Putamen_R' | 4.44235292 | 0.00015813 |
| 'Frontal_Inf_Tri_L' | 4.492214975 | 0.000138966 | 'Temporal_Pole_Mid_R' | 2.376185441 | 0.02816383 | 'Frontal_Inf_Tri_L' | 4.492214975 | 0.000138966 | 'Frontal_Sup_Orb_R' | 4.48960514 | 0.00013991 |
| 'Temporal_Sup_R' | 4.496621545 | 0.000137388 | 'Temporal_Pole_Mid_L' | 2.391370397 | 0.027290219 | 'Temporal_Sup_R' | 4.496621545 | 0.000137388 | 'Insula_L' | 4.51026584 | 0.00013261 |
| 'Frontal_Inf_Orb_R' | 4.516437121 | 0.000130509 | 'Parietal_Sup_L' | 2.391822057 | 0.027264625 | 'Frontal_Inf_Orb_R' | 4.516437121 | 0.000130509 | 'Supp_Motor_Area_L' | 4.52492064 | 0.00012767 |
| 'Frontal_Mid_R' | 4.516565998 | 0.000130466 | 'Lingual_L' | 2.400696777 | 0.02676624 | 'Frontal_Mid_R' | 4.516565998 | 0.000130466 | 'Paracentral_Lobule_R' | 4.56903088 | 0.00011387 |
| 'Parietal_Inf_L' | 4.559426357 | 0.000116745 | 'Parietal_Sup_R' | 2.473592944 | 0.022982791 | 'Parietal_Inf_L' | 4.559426357 | 0.000116745 | 'Insula_R' | 4.59918828 | 0.00010531 |
| 'Frontal_Inf_Oper_R' | 4.613938514 | 0.000101357 | 'ParaHippocampal_R' | 2.488600592 | 0.022268783 | 'Frontal_Inf_Oper_R' | 4.613938514 | 0.000101357 | 'Caudate_L' | 4.63646555 | 9.56E-05 |
| 'Frontal_Mid_L' | 4.631376218 | 9.69E-05 | 'Insula_R' | 2.524087026 | 0.020662455 | 'Frontal_Mid_L' | 4.631376218 | 9.69E-05 | 'Precentral_L' | 4.64770095 | 9.29E-05 |
| 'Rolandic_Oper_L' | 4.637847806 | 9.53E-05 | 'Frontal_Inf_Orb_R' | 2.524769907 | 0.02063264 | 'Rolandic_Oper_L' | 4.637847806 | 9.53E-05 | 'Frontal_Inf_Tri_R' | 4.66970797 | 8.77E-05 |
| 'Parietal_Sup_R' | 4.63900656 | 9.50E-05 | 'Frontal_Inf_Oper_L' | 2.534687926 | 0.020204152 | 'Parietal_Sup_R' | 4.63900656 | 9.50E-05 | 'Precuneus_L' | 4.83177132 | 5.76E-05 |
| 'Pallidum_R' | 4.646147343 | 9.32E-05 | 'Calcarine_R' | 2.543595967 | 0.019826454 | 'Pallidum_R' | 4.646147343 | 9.32E-05 | 'Frontal_Sup_Medial_L' | 4.87134209 | 5.20E-05 |
| 'Amygdala_L' | 4.683076202 | 8.47E-05 | 'Frontal_Inf_Tri_L' | 2.572095615 | 0.0186623 | 'Amygdala_L' | 4.683076202 | 8.47E-05 | 'Temporal_Mid_L' | 4.92461292 | 4.53E-05 |
| 'Frontal_Sup_L' | 4.741312461 | 7.28E-05 | 'Precuneus_L' | 2.627996642 | 0.01656387 | 'Frontal_Sup_L' | 4.741312461 | 7.28E-05 | 'Occipital_Sup_R' | 4.93179445 | 4.45E-05 |
| 'Cingulum_Mid_L' | 4.767266776 | 6.81E-05 | 'Frontal_Inf_Orb_L' | 2.641576925 | 0.016088935 | 'Cingulum_Mid_L' | 4.767266776 | 6.81E-05 | 'Frontal_Sup_Medial_R' | 4.93903845 | 4.36E-05 |
| 'Frontal_Sup_R' | 4.822792353 | 5.90E-05 | 'Heschl_R' | 2.648788494 | 0.015842009 | 'Frontal_Sup_R' | 4.822792353 | 5.90E-05 | 'Calcarine_R' | 4.94237421 | 4.33E-05 |
| 'Temporal_Mid_L' | 4.833898458 | 5.73E-05 | 'Insula_L' | 2.701298676 | 0.014149583 | 'Temporal_Mid_L' | 4.833898458 | 5.73E-05 | 'Frontal_Inf_Tri_L' | 4.97866883 | 3.94E-05 |
| 'Putamen_L' | 4.861224844 | 5.34E-05 | 'Parietal_Inf_L' | 2.717073499 | 0.013675734 | 'Putamen_L' | 4.861224844 | 5.34E-05 | 'Frontal_Sup_L' | 4.98480678 | 3.88E-05 |
| 'Frontal_Inf_Tri_R' | 4.864014175 | 5.30E-05 | 'Temporal_Pole_Sup_L' | 2.745372292 | 0.012863353 | 'Frontal_Inf_Tri_R' | 4.864014175 | 5.30E-05 | 'Postcentral_R' | 5.05830794 | 3.21E-05 |
| 'Putamen_R' | 4.943414531 | 4.31E-05 | 'Frontal_Mid_Orb_R' | 2.777862232 | 0.011987475 | 'Putamen_R' | 4.943414531 | 4.31E-05 | 'Parietal_Inf_L' | 5.06612368 | 3.14E-05 |
| 'Heschl_R' | 4.995347196 | 3.77E-05 | 'Supp_Motor_Area_L' | 2.797734025 | 0.011480191 | 'Heschl_R' | 4.995347196 | 3.77E-05 | 'Parietal_Sup_L' | 5.07101403 | 3.10E-05 |

| | | | | | | | | | | | |
|---|---|---|---|---|---|---|---|---|---|---|---|
| 'Frontal_Mid_Orb_R' | 5.066433933 | 3.14E-05 | 'Cingulum_Mid_R' | 2.814822303 | 0.011060463 | 'Frontal_Mid_Orb_R' | 5.066433933 | 3.14E-05 | 'Rectus_R' | 5.12582969 | 2.69E-05 |
| 'Temporal_Sup_L' | 5.083784156 | 3.00E-05 | 'Rolandic_Oper_L' | 2.856740858 | 0.010092257 | 'Temporal_Sup_L' | 5.083784156 | 3.00E-05 | 'Paracentral_Lobule_L' | 5.20695672 | 2.18E-05 |
| 'Frontal_Med_Orb_R' | 5.092987422 | 2.93E-05 | 'Parietal_Inf_R' | 2.91978164 | 0.008787964 | 'Frontal_Med_Orb_R' | 5.092987422 | 2.93E-05 | 'Frontal_Inf_Orb_R' | 5.21022028 | 2.17E-05 |
| 'Heschl_L' | 5.130795552 | 2.66E-05 | 'Frontal_Med_Orb_L' | 2.944111592 | 0.008329357 | 'Heschl_L' | 5.130795552 | 2.66E-05 | 'Thalamus_L' | 5.25070693 | 1.95E-05 |
| 'Cingulum_Mid_R' | 5.133398599 | 2.64E-05 | 'Frontal_Sup_R' | 3.001723707 | 0.00733368 | 'Cingulum_Mid_R' | 5.133398599 | 2.64E-05 | 'Precuneus_R' | 5.32066865 | 1.63E-05 |
| 'Frontal_Sup_Orb_L' | 5.155353638 | 2.49E-05 | 'Frontal_Mid_L' | 3.043047263 | 0.0066915 | 'Frontal_Sup_Orb_L' | 5.155353638 | 2.49E-05 | 'Thalamus_R' | 5.51421221 | 9.91E-06 |
| 'Angular_R' | 5.238751277 | 2.01E-05 | 'Heschl_L' | 3.049254423 | 0.006599875 | 'Angular_R' | 5.238751277 | 2.01E-05 | 'Rolandic_Oper_L' | 5.60666974 | 7.83E-06 |
| 'Pallidum_L' | 5.259328595 | 1.91E-05 | 'Temporal_Pole_Sup_R' | 3.083511284 | 0.006115693 | 'Pallidum_L' | 5.259328595 | 1.91E-05 | 'Cingulum_Mid_L' | 5.6276725 | 7.42E-06 |
| 'Frontal_Sup_Orb_R' | 5.337381754 | 1.56E-05 | 'Cingulum_Mid_L' | 3.14961011 | 0.005277132 | 'Frontal_Sup_Orb_R' | 5.337381754 | 1.56E-05 | 'Frontal_Sup_Orb_L' | 5.63888593 | 7.21E-06 |
| 'Calcarine_R' | 5.387129282 | 1.37E-05 | 'Olfactory_L' | 3.254365215 | 0.004172579 | 'Calcarine_R' | 5.387129282 | 1.37E-05 | 'Supp_Motor_Area_R' | 5.74302834 | 5.53E-06 |
| 'Thalamus_L' | 5.395038908 | 1.35E-05 | 'Precentral_R' | 3.315737081 | 0.00363416 | 'Thalamus_L' | 5.395038908 | 1.35E-05 | 'Frontal_Mid_R' | 5.7794818 | 5.04E-06 |
| 'Insula_L' | 5.413356041 | 1.28E-05 | 'Occipital_Mid_R' | 3.418194809 | 0.002883292 | 'Insula_L' | 5.413356041 | 1.28E-05 | 'Rectus_L' | 5.78703859 | 4.94E-06 |
| 'Olfactory_L' | 5.443556517 | 1.19E-05 | 'Precentral_L' | 3.428592842 | 0.002816216 | 'Olfactory_L' | 5.443556517 | 1.19E-05 | 'Cingulum_Ant_R' | 5.80705259 | 4.70E-06 |
| 'Paracentral_Lobule_R' | 5.473420237 | 1.10E-05 | 'Rectus_R' | 3.47985 | 0.002507387 | 'Paracentral_Lobule_R' | 5.473420237 | 1.10E-05 | 'Putamen_L' | 5.94164701 | 3.34E-06 |
| 'Parietal_Inf_R' | 5.488782585 | 1.06E-05 | 'Cingulum_Ant_L' | 3.610253273 | 0.001864437 | 'Parietal_Inf_R' | 5.488782585 | 1.06E-05 | 'Pallidum_L' | 5.9717362 | 3.10E-06 |
| 'Frontal_Sup_Medial_L' | 5.497743621 | 1.03E-05 | 'Pallidum_R' | 3.649313268 | 0.0017058 | 'Frontal_Sup_Medial_L' | 5.497743621 | 1.03E-05 | 'Frontal_Sup_R' | 6.06915617 | 2.42E-06 |
| 'Parietal_Sup_L' | 5.544786211 | 9.17E-06 | 'Frontal_Inf_Oper_R' | 3.652156311 | 0.00169479 | 'Parietal_Sup_L' | 5.544786211 | 9.17E-06 | 'Frontal_Med_Orb_L' | 6.19995703 | 1.75E-06 |
| 'Caudate_L' | 5.686636907 | 6.38E-06 | 'Thalamus_R' | 3.791884994 | 0.001232388 | 'Caudate_L' | 5.686636907 | 6.38E-06 | 'Occipital_Mid_L' | 6.24209067 | 1.57E-06 |
| 'Angular_L' | 5.701356273 | 6.15E-06 | 'Frontal_Sup_Orb_L' | 3.87616798 | 0.001016667 | 'Angular_L' | 5.701356273 | 6.15E-06 | 'Cingulum_Mid_R' | 6.53653119 | 7.58E-07 |
| 'Frontal_Med_Orb_L' | 5.751999993 | 5.40E-06 | 'Frontal_Sup_L' | 4.199710844 | 0.000485637 | 'Frontal_Med_Orb_L' | 5.751999993 | 5.40E-06 | 'Frontal_Med_Orb_R' | 6.58760157 | 6.68E-07 |
| 'Rectus_R' | 5.808588701 | 4.68E-06 | 'Frontal_Sup_Medial_R' | 4.397815144 | 0.000309252 | 'Rectus_R' | 5.808588701 | 4.68E-06 | 'Cingulum_Ant_L' | 6.70893508 | 4.96E-07 |
| 'Cingulum_Ant_L' | 6.37957974 | 1.12E-06 | 'Supp_Motor_Area_R' | 5.499349631 | 2.64E-05 | 'Cingulum_Ant_L' | 6.37957974 | 1.12E-06 | 'Frontal_Mid_L' | 6.79494914 | 4.02E-07 |

**Supplementary Table 3:** Region wise changes in PILI for MCS and UWS patients after simulating LSD and psilocybin. Assessed by paired T-tests.

#### Baseline structural and functional differences between patients with DoC and their correlations with simulated treatment effects

| Network | F statistic | ANOVA p-Value | Post-hoc Comparison | Mean Difference | p-Value (Post-hoc) |
|---|---|---|---|---|---|
| Mean FC | 4.493 | 0.0142 | MCS vs. UWS | -3.491 | 0.419 |
| | | | MCS vs. CNT | -11.106 | 0.202 |
| | | | UWS vs. CNT | -16.560 | 0.012 |
| Attention | 5.071 | 0.0433 | MCS vs. UWS | 0.0567 | 0.4538 |
| | | | MCS vs. CNT | -0.0554 | 0.3711 |
| | | | UWS vs. CNT | -0.1122 | 0.0356 |
| Auditory | 13.232 | <0.001 | MCS vs. UWS | 0.0847 | 0.1674 |
| | | | MCS vs. CNT | -0.1086 | 0.0234 |
| | | | UWS vs. CNT | -0.1934 | 0.0001 |
| Default Mode | 7.177 | 0.0082 | MCS vs. UWS | 0.0934 | 0.0658 |
| | | | MCS vs. CNT | -0.0291 | 0.6952 |
| | | | UWS vs. CNT | -0.1225 | 0.0062 |
| Frontoparietal | 6.343 | 0.0088 | MCS vs. UWS | 0.0207 | 0.8637 |
| | | | MCS vs. CNT | -0.0856 | 0.0427 |
| | | | UWS vs. CNT | -0.1062 | 0.0168 |
| Limbic | 6.039 | 0.0139 | MCS vs. UWS | 0.0658 | 0.2917 |
| | | | MCS vs. CNT | -0.0569 | 0.2974 |
| | | | UWS vs. CNT | -0.1227 | 0.0104 |
| Precuneus | 10.076 | 0.0014 | MCS vs. UWS | 0.0558 | 0.3229 |
| | | | MCS vs. CNT | -0.0774 | 0.0612 |
| | | | UWS vs. CNT | -0.1333 | 0.0013 |
| Sensorimotor | 6.899 | 0.0123 | MCS vs. UWS | 0.0585 | 0.4098 |
| | | | MCS vs. CNT | -0.07 | 0.1899 |
| | | | UWS vs. CNT | -0.1285 | 0.0105 |
| Thalamus | 10.496 | <0.001 | MCS vs. UWS | 0.0652 | 0.2111 |
| | | | MCS vs. CNT | -0.0823 | 0.0414 |
| | | | UWS vs. CNT | -0.1474 | 0.0003 |
| Visual | 13.232 | <0.001 | MCS vs. UWS | 0.0541 | 0.288 |
| | | | MCS vs. CNT | -0.1211 | 0.0006 |
| | | | UWS vs. CNT | -0.1752 | <0.001 |

**Supplementary Table 4:** Results of one-way ANOVA of functional connectivity in 9 resting state networks and the overall average FC in overall brain regions between MCS and UWS patients and healthy controls.

| Metric | F-statistic | ANOVA p-Value | Post-hoc Comparison | Mean Difference | p-Value (Post-hoc) |
|---|---|---|---|---|---|
| Mean SC | 1.294 | 0.1524 | MCS vs. UWS | 0.0225 | 0.4114 |
| | | | MCS vs. CNT | -0.0106 | 0.7833 |
| | | | UWS vs. CNT | -0.0331 | 0.1303 |
| Graph strength | 6.305 | 0.0028 | MCS vs. UWS | 0.0339 | 0.1207 |
| | | | MCS vs. CNT | -0.0242 | 0.2596 |
| | | | UWS vs. CNT | -0.0581 | 0.0018 |
| Global efficiency | 4.819 | 0.0104 | MCS vs. UWS | 0.0014 | 0.1994 |
| | | | MCS vs. CNT | -0.001 | 0.3497 |
| | | | UWS vs. CNT | -0.0023 | 0.0073 |
| Local efficiency | 2.390 | 0.0978 | MCS vs. UWS | -0.0001 | 0.6021 |
| | | | MCS vs. CNT | -0.0002 | 0.08 |
| | | | UWS vs. CNT | -0.0001 | 0.5538 |
| Centrality | 5.105 | 0.0081 | MCS vs. UWS | -9.916 | 0.069 |
| | | | MCS vs. CNT | 3.4527 | 0.6602 |
| | | | UWS vs. CNT | 13.3687 | 0.0063 |
| Characteristic Path length | 39.149 | <0.001 | MCS vs. UWS | -1.4957 | <0.001 |
| | | | MCS vs. CNT | 1.2375 | <0.001 |
| | | | UWS vs. CNT | 2.7332 | <0.001 |
| Fractional anisotropy | 67.491 | <0.001 | MCS vs. UWS | 0.012 | 0.1204 |
| | | | MCS vs. CNT | -0.0486 | <0.001 |
| | | | UWS vs. CNT | -0.0606 | <0.001 |

**Supplementary Table 5:** Results of one-way ANOVA of structural connectivity graph metrics and the overall average structural connectivity over all brain regions between MCS and UWS patients and healthy controls.

| Group | Condition | Centrality | Characteristic path length | Fractional anisotropy | Global efficiency | Local efficiency | Graph strength |
|---|---|---|---|---|---|---|---|
| UWS + LSD | R-value | -0.524 | -0.384 | 0.567 | 0.541 | -0.088 | 0.534 |
| | P-value | 0.018 | 0.095 | 0.009 | 0.014 | 0.711 | 0.015 |
| UWS + psilocybin | R-value | -0.328 | -0.382 | 0.388 | 0.371 | 0.494 | 0.493 |
| | P-value | 0.158 | 0.096 | 0.091 | 0.107 | 0.027 | 0.027 |
| MCS + LSD | R-value | -0.260 | -0.124 | -0.150 | 0.688 | -0.033 | 0.095 |
| | P-value | 0.200 | 0.547 | 0.462 | 0.139 | 0.872 | 0.645 |
| MCS + psilocybin | R-value | -0.396 | 0.309 | -0.186 | -0.111 | -0.140 | -0.098 |
| | P-value | 0.045 | 0.124 | 0.363 | 0.591 | 0.494 | 0.633 |

**Supplementary Table 6:** Results of correlational analyses between baseline structural graph theory metrics and the changes in PILI values as a result of simulating LSD and psilocybin in patients with UWS and patients in the MCS.

#### Supplementary analyses

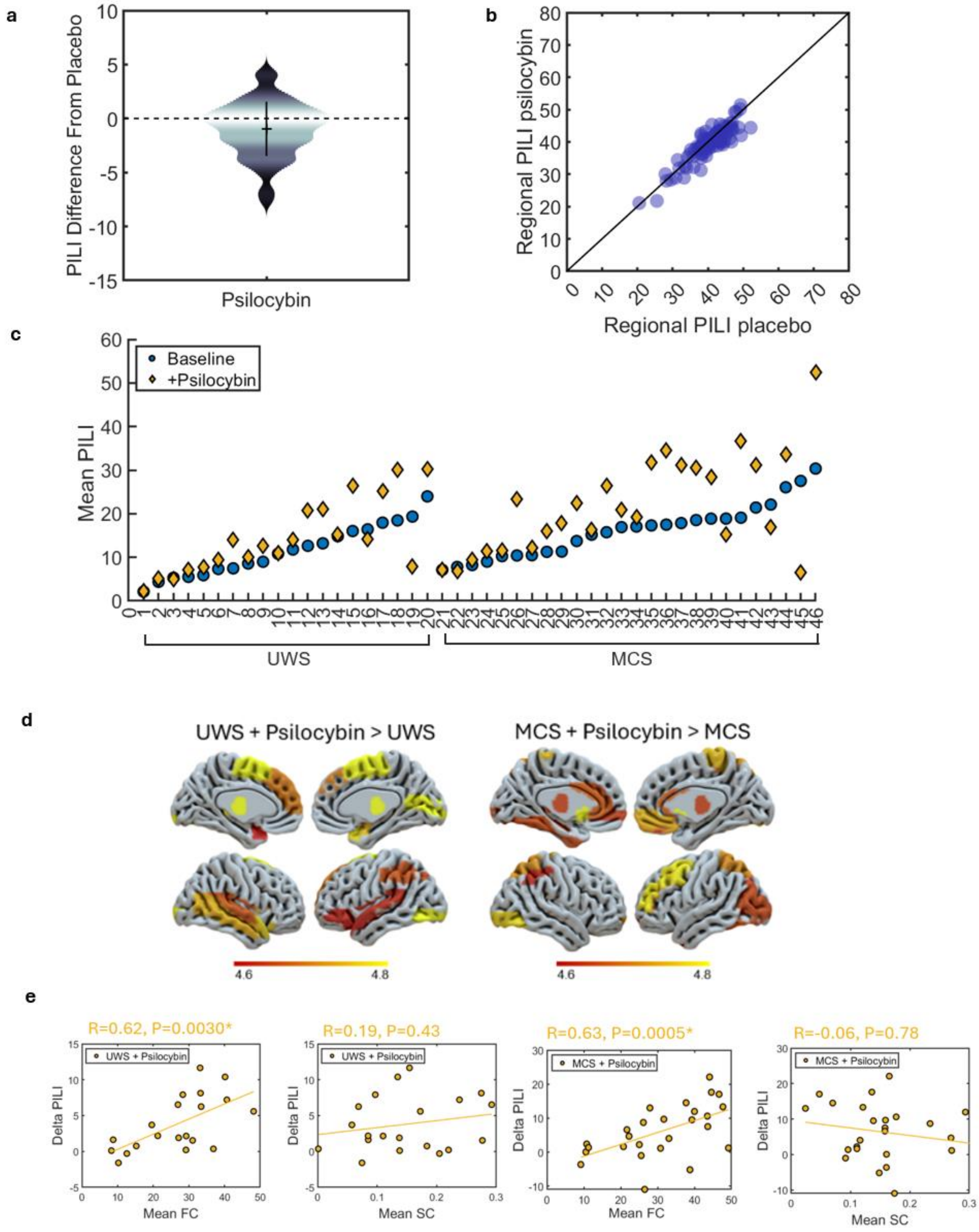

**Supplementary Figure 2:** Supplementary analysis using another psilocybin dataset with a between-subject's design. **a.** Violin plot of the distribution of PILI trials averaged over all brain regions of the group level model of psilocybin. Y axis represents the distance of PILI values in the psilocybin condition compared to the placebo condition. **b.** Absolute region wise PILI in the psilocybin condition X axis plotted against the placebo condition Y axis. **c.** Individual patient models before and after simulation of psilocybin. Blue circles are each patient at baseline, yellow diamonds represent the patient after the simulation of psilocybin. **d.** Results of region wise t-tests displaying brain regions with significant increases, in PILI from simulating psilocybin in patients with UWS (left) and MCS patients (right), Bonferroni corrected for multiple comparisons for the 90 brain regions. **e.** Scatter plots showing correlations between the changing in PILI values as a result of simulating psilocybin (Delta PILI) and the average baseline SC and FC in UWS patients (Left) and MCS patients (Right). Regression lines are indicated in yellow. Associated correlation coefficients (R) and p-values are stated above, star indicates significance, after Bonferroni correction or the 4 comparisons.

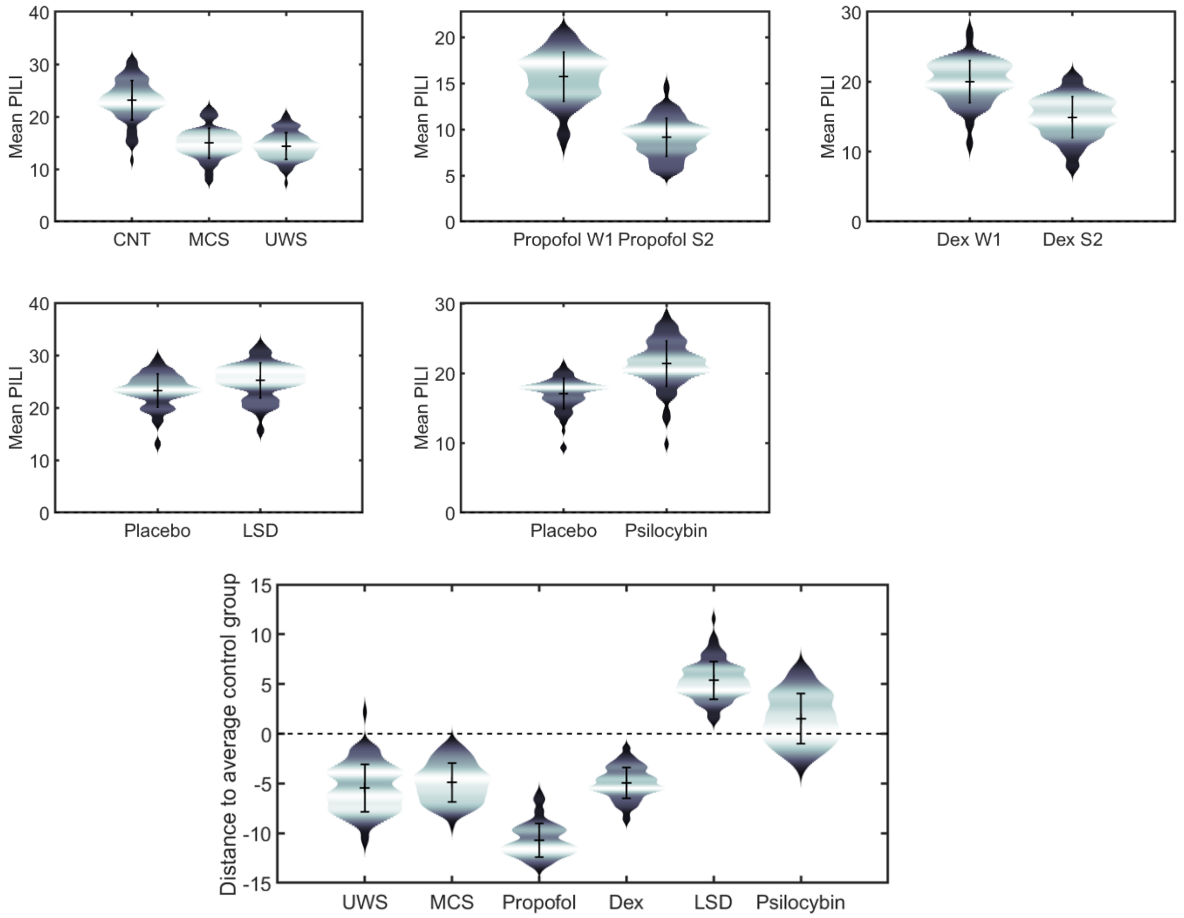

**Supplementary Figure 3:** a. Violin plots of the distribution of PILI trials averaged over all brain regions of group level models in each state of consciousness. Top. Absolute PILI values in each state of consciousness and the respective comparison condition. Bottom. PILI values in each state of consciousness given in reference to the distance of each group from the average of all the comparison groups: healthy controls for the UWS and MCS groups, the respective wakefulness conditions for propofol and dexmedetomidine, respective placebo conditions for LSD and psilocybin.

**Supplementary Discussion 2:** The differences between acquisition protocols and scanning parameters, mean that comparing PILI values between datasets is challenging. This is evident from the variability in PILI values across different control group models (Figure S4). Despite this variability, using the average across control/comparison groups as a reference for each condition mostly maintains the relationships within and between datasets. However, according to these results it would suggest that propofol anaesthesia has the least critical brain dynamics, which conflicts with previous work. These factors highlight the robustness of the method within some bounds at the group level, but also exemplify its limitations and the unsuitability in generalising between datasets. These results are likely caused by the inherent variability in PILI between subjects. The supplementary analyses of psilocybin with a between-subject design further support this idea. The PILI values in the psilocybin group compared to the placebo group showed a slight deviation towards decreases in the psilocybin group, if not no difference (Cohens d = -0.029) (Figure S5a). Similarly, the regional distribution of PILI values indicates that most values tend to cluster around the zero-line or a slight negative deviation, indicating no difference between conditions (Figure S5b). Critically, the psilocybin Maastricht dataset employs a between-subjects design. This means that when comparing PILI values between conditions, the different subjects who received psilocybin may have different baseline PILI values. To further illustrate this, we observe that PILI can vary across single-subject models of healthy controls, with values ranging from 10 to 30 (Figure S1b). Therefore, the naturally occurring range of PILI values at wakefulness between the two groups (i.e., those receiving psilocybin and those receiving placebo) may have influenced the absence of increases in PILI seen in this dataset. This highlights the point that whilst PILI can discriminate between states of consciousness, it should ideally only be used in a within-subjects design. Furthermore, when estimating the treatment effect, using different subjects for each condition means that both structural and functional connectivity changed between conditions. In a within-subject design, only functional connectivity changes as a result of drug administration. Although we can generally assume that within some bounds, all healthy subjects have similar structural connectivity, the specificity of the virtual pharmacology method, which uses changes in local bifurcation parameters to estimate changes in brain functional dynamics likely benefits from estimating this change within the same subjects, thus keeping the structure constant. This suboptimal estimation of treatment effects for this specific use case likely underlies the absence of correlation between the baseline structural connectivity and changes in PILI as a result of simulating psilocybin in patients with UWS which were strongly correlated in the other two psychedelic datasets. Therefore, this dataset demonstrates the need for subject specificity both in comparing the brain dynamics between conditions and when estimating the treatment effect through extracting parameter sets.
